# OMIO: A policy-driven Python library for reproducible microscopy image I/O

**DOI:** 10.64898/2026.06.09.731118

**Authors:** Fabrizio Musacchio, Henrike Antony, Arush Baijal, Sophie Crux, Falko Fuhrmann, Nala Gockel, Denise Marie Hoffmann, Dilek Mercan, Felix Christopher Nebeling, Katharina Wolff, Martin Fuhrmann

## Abstract

Modern fluorescence and multiphoton microscopy workflows operate within a heterogeneous ecosystem of file formats, partially overlapping metadata standards, and reader-specific conventions. In practice, this frequently leads to silent axis misinterpretations, loss or corruption of physical voxel size information, and laboratory-specific glue code that is fragile, poorly documented, and difficult to reproduce. OMIO, short for *Open Microscopy Image I/O*, addresses these issues by providing a lightweight, policy-driven image I/O layer for Python that enforces a canonical, OME-compatible data representation at the API boundary. The central contribution of OMIO is the explicit separation of low-level format access from semantic normalization. Existing reader libraries are used as interchangeable backends for extracting pixel data and available metadata, while OMIO enforces axis conventions, metadata interpretation, and fallback decisions in a centralized and auditable policy layer. This design allows heterogeneous microscopy inputs to be converted into a stable representation without propagating backend-specific assumptions into downstream analysis code.

The core design principles of OMIO include canonical axis semantics (TZCYX), robust metadata normalization with explicit and auditable fallbacks, memory-aware operation via optional Zarr-based backends, and workflow-level semantics that extend beyond individual files to folder stacks and BIDS-like project structures. This architecture allows OMIO to orchestrate existing reader libraries into a coherent and reproducible I/O pipeline without replacing or duplicating their functionality.

OMIO is implemented as an open-source and community-oriented system in which support for additional file formats and metadata conventions can be added incrementally through modular reader backends. By encouraging the contribution of example datasets, backend extensions, and feature requests, OMIO is designed to evolve alongside emerging acquisition systems while preserving strict semantic guarantees at the interface level. The resulting standardized OME-TIFF outputs are immediately suitable for downstream quantitative analysis and interactive inspection in scientific Python workflows, including workflows based on ImageJ and Napari. By standardizing image data at the I/O boundary, OMIO supports FAIR-aligned data sharing and reproducible microscopy analysis while facilitating the development of interoperable downstream tools.

## Introduction

Quantitative fluorescence and multiphoton microscopy critically depend on the correct interpretation of image axes, physical voxel sizes, and acquisition metadata. Errors in axis ordering or pixel spacing directly propagate into downstream measurements, affecting estimates of distances, volumes, motility metrics, growth rates, and temporal dynamics. Crucially, such errors are often silent: numerically plausible results may be produced that are nevertheless scientifically invalid, and may remain undetected throughout an analysis pipeline[1, 2].

To address interoperability and metadata consistency, the Open Microscopy Environment introduced the OME data model and OME-TIFF as a standardized container for multidimensional image data and associated metadata[3]. The OME model provides a formal specification for axis semantics, physical units, and acquisition context, and has been widely adopted by imaging software, repositories, and analysis tools. Nevertheless, in practical microscopy workflows, data ingestion remains fragmented across file formats, vendor-specific metadata conventions, and heterogeneous reader implementations. As a result, identical datasets may be interpreted differently depending on the reader, software version, or downstream analysis environment.

In everyday laboratory practice, microscopy data ingestion and conversion are frequently handled by ad hoc scripts or format-specific utilities. Such workflows often make implicit assumptions about the order and meaning of image axes. They may also remove singleton dimensions, that is, axes of length one, overwrite metadata fields, or apply undocumented fallback rules when required information is missing or ambiguous. While numerous libraries exist that provide access to individual microscopy formats, they typically expose low-level reader outputs and delegate semantic interpretation to downstream code. As a consequence, the burden of ensuring consistent axis semantics and metadata integrity is shifted to analysis pipelines, where errors become harder to detect, correct, and document.

Across formats, tools, and workflows, these challenges manifest in a small number of recurring and systematic failure patterns in microscopy data ingestion[1, 2]. For clarity, these patterns are summarized as distinct failure modes in Box 1. They are not isolated edge cases, but structural issues that arise in heterogeneous and long-lived microscopy projects and that directly motivate the need for explicit semantic policies at the point of data ingestion.

One prominent example is axis ambiguity and silent reordering across file formats and reader implementations, which can lead to misinterpretation of spatial, temporal, or channel dimensions without triggering explicit errors (Failure mode 1 in Box 1). Even when a formal standard such as OME-TIFF is used, implicit squeezing of singleton dimensions and shape-dependent logic can reintroduce ambiguity at the interface between readers and analysis code (Failure mode 2 in Box 1).

Physical voxel sizes represent another critical point of failure. They are essential for quantitative interpretation, yet are frequently missing, incomplete, or inconsistently encoded across formats[4]. In many workflows, missing voxel sizes are silently replaced by defaults or ignored altogether, directly compromising measurements while remaining numerically plausible (Failure mode 3 in Box 1). Additional failure modes arise from undocumented, format-specific metadata heuristics that may change across library versions (Failure mode 4), as well as from workflow fragmentation across graphical applications, conversion utilities, and scripting environments, each of which may reinterpret or discard metadata differently (Failure mode 5 in Box 1). These issues are further amplified in batch processing scenarios, where custom scripts may apply inconsistent logic across datasets processed within the same study (Failure mode 6 in Box 1).

Taken together, these observations reveal a conceptual gap between file format support and reproducible scientific analysis. Format readers and visualization tools alone do not guarantee semantic consistency across datasets, especially when workflows involve multiple acquisition systems, legacy data, or evolving software environments. What is often missing is an explicit and enforceable policy layer that defines how image data and metadata are interpreted, normalized, and validated at the point of ingestion, and that does so uniformly across formats and workflows.

OMIO, short for *Open Microscopy Image I/O*, is designed to address this gap by acting as a policy-driven image I/O layer on top of existing microscopy readers. Rather than aiming for maximal format coverage, OMIO enforces a canonical, OME-compatible data representation at the API boundary. Axis semantics are made explicit and normalized to a single convention, singleton dimensions are preserved rather than implicitly removed, physical voxel sizes are treated as first-class metadata, and all fallback decisions are surfaced to the user. In doing so, OMIO directly targets the failure modes summarized in Box 1 by shifting critical semantic decisions upstream into a transparent and inspectable I/O layer.

In this sense, OMIO is not merely a file reader or converter, but an infrastructure component that promotes reproducibility by construction. It formalizes common but often implicit assumptions about microscopy data into explicit policies, providing a stable and auditable foundation for downstream quantitative analysis across heterogeneous datasets and computational environments[5].

#### Box 1: Failure modes in practical microscopy data ingestion

Despite the availability of standardized file formats and mature analysis software, several recurring failure modes are commonly observed in practical microscopy workflows.

**Failure mode 1: Axis ambiguity and silent reordering.**

Microscopy file formats frequently store image data in backend-specific axis orders. Reader libraries may return arrays in different conventions depending on format, acquisition settings, or library versions. Downstream analysis code often assumes a fixed axis order, leading to silent misinterpretation of spatial, temporal, or channel dimensions without triggering explicit errors.

**Failure mode 2: Implicit dimension squeezing and shape-dependent logic.**

Singleton dimensions, such as time or channel axes of length one, are often dropped implicitly by readers or analysis code. This leads to shape-dependent branching in downstream pipelines and breaks assumptions when datasets with different dimensionality are processed together.

**Failure mode 3: Loss or corruption of physical pixel sizes.**

Physical voxel sizes are essential for quantitative measurements, yet they are frequently missing, incomplete, or inconsistently encoded across formats. In many workflows, missing voxel sizes are silently replaced by default values or ignored entirely, resulting in incorrect physical measurements that remain numerically plausible.

**Failure mode 4: Format-specific metadata heuristics.**

Reader implementations often apply undocumented heuristics when interpreting metadata fields, particularly for legacy or vendor-specific formats. These heuristics may change across library versions, undermining long-term reproducibility and comparability of results.

**Failure mode 5: Workflow fragmentation across tools.**

Data are commonly inspected in graphical applications, converted via standalone utilities, and analyzed in scripting environments. Each step may reinterpret or discard metadata differently, making it difficult to ensure that the data entering an analysis pipeline faithfully represent the original acquisition.

**Failure mode 6: Batch processing inconsistencies.**

Batch processing of microscopy datasets often involves custom scripts that apply format-specific logic. Without a unified policy layer, these scripts may introduce inconsistencies in axis handling or metadata interpretation across datasets, leading to heterogeneous analysis inputs.

These failure modes are not isolated edge cases but arise systematically in heterogeneous and long-lived microscopy projects. Addressing them requires explicit and enforceable semantic policies at the point of data ingestion.

## Related work

Microscopy image data are supported by a wide ecosystem of software libraries, end-user applications, and file format standards. These tools address complementary aspects of image acquisition, visualization, and analysis, but they differ substantially in how explicitly they handle axis semantics, metadata normalization, and reproducibility across heterogeneous workflows. In particular, many existing tools leave critical semantic decisions implicit, thereby exposing users to several of the recurring failure modes summarized in Box 1.

### Reader libraries and format abstractions

Several libraries provide programmatic access to microscopy image formats. The Python library *tifffile*[6] offers comprehensive support for TIFF-based formats, including OMETIFF, and exposes low-level access to image arrays and embedded metadata. Similarly, the Bio-Formats[7] library provides broad format coverage through a Java-based framework and serves as the backbone for many desktop microscopy applications. More recently, *aicsimageio*[8] has aimed to provide a unified Python interface to selected microscopy formats by abstracting over multiple backends.

While these libraries are indispensable for format access, they primarily expose reader-specific representations of image data and metadata. Axis ordering, dimensionality, and physical pixel sizes are often returned in backend-dependent forms, and semantic normalization is left to downstream code. As a result, analysis pipelines built directly on top of reader libraries frequently reimplement axis handling and metadata interpretation logic in an ad hoc and non-reproducible manner, directly reflecting Failure modes 1 and 2 in Box 1. Physical voxel sizes may be inconsistently exposed or partially missing, placing the responsibility for correct quantitative interpretation on downstream analysis code (Failure mode 3).

Vendor-specific utilities such as *czifile*[9] or *utils2p*[10] address individual proprietary formats but similarly focus on data extraction rather than semantic unification. In particular, they tend to mirror the structure of the underlying file format, which may not align with analysis-oriented representations or cross-format consistency requirements. Metadata interpretation in these tools often relies on format-specific heuristics that are undocumented and subject to change, thereby exacerbating reproducibility issues over time (Failure mode 4 in Box 1).

In Table 1, we summarize key features of several popular microscopy I/O libraries, highlighting their capabilities and limitations with respect to format support, metadata handling, large data support, workflow-level semantics, output guarantees, and project characteristics.

**Table 1:**
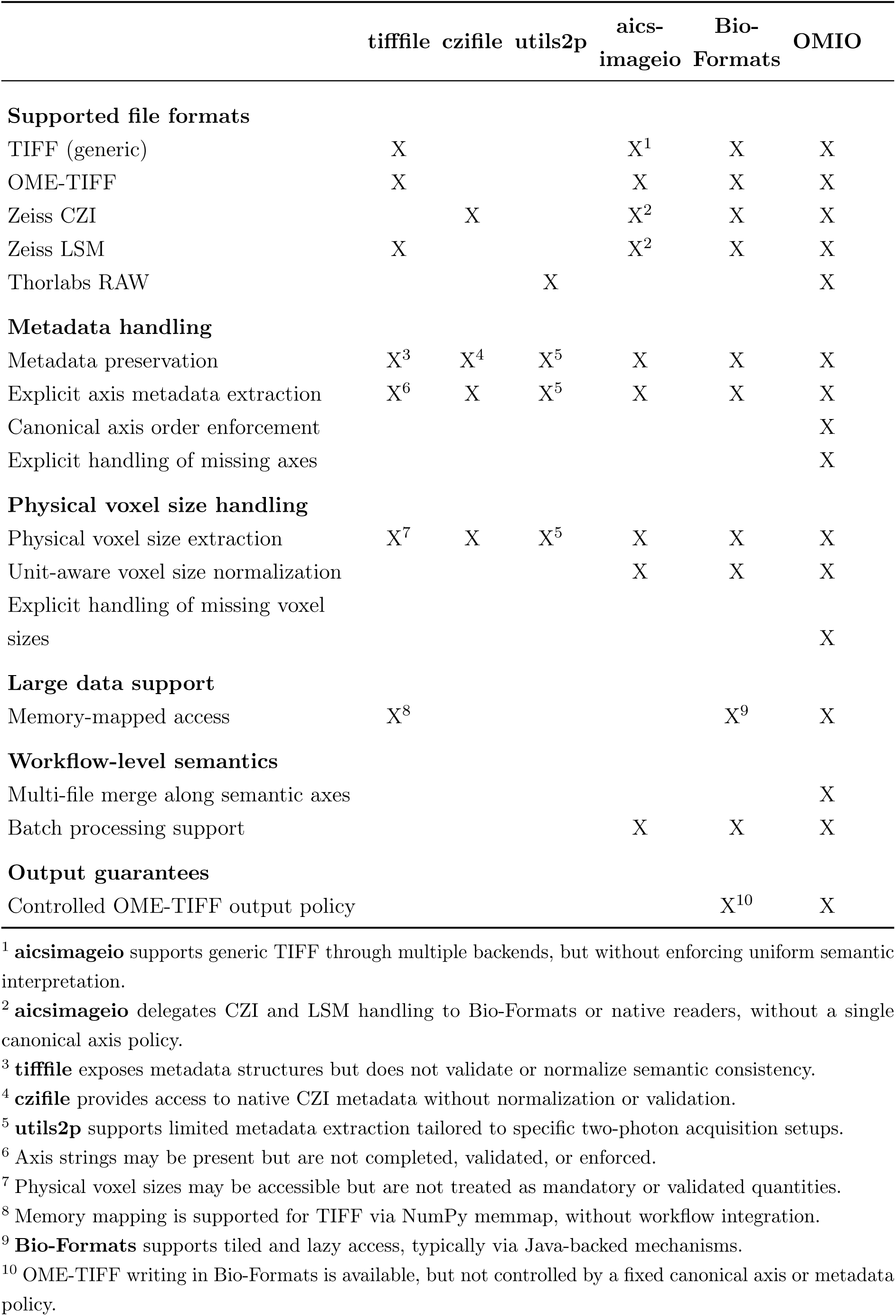
Feature comparison of microscopy I/O libraries. The table summarizes key capabilities related to format support, metadata handling, large data support, workflow-level semantics, output guarantees, and project characteristics across several popular open source microscopy I/O libraries. An extended comparison is provided in Table 3.

### End-user applications and visualization platforms

Graphical applications such as ImageJ/Fiji[11, 12], Napari[13, 14], Imaris[15], and vendor software suites including Zeiss ZEN[16] or Leica LAS X[17] provide powerful tools for visualization, annotation, and interactive analysis. These platforms are highly effective for exploratory work and manual inspection, and many of them internally support OME-compatible metadata models[3].

However, these applications are not designed to act as explicit policy layers for data ingestion in automated or reproducible pipelines. Axis interpretation and metadata handling are typically embedded implicitly within the application logic and user interface, rather than being surfaced as programmable and inspectable decisions. When data are exported or transferred between platforms, semantic assumptions may be flattened, reordered, or partially lost, particularly when workflows involve batch processing, custom analysis scripts, or long-term data reuse. These characteristics directly relate to Failure modes 1, 2, and 5 in Box 1.

Moreover, while desktop applications provide internally consistent semantics within their own environment, they do not address the problem of enforcing a canonical representation across heterogeneous computational workflows. For example, the same dataset may be interpreted differently when opened in Fiji, Napari, or a Python-based analysis pipeline, depending on how axes and metadata are mapped internally. Such discrepancies are especially problematic when datasets are processed programmatically or reused across projects, where implicit application-specific assumptions are no longer visible.

### Conceptual gap between access and policy

Taken together, existing tools largely separate into two categories: low-level readers that provide access to image data without enforcing semantic consistency, and highlevel applications that embed semantic decisions implicitly but do not expose them as reproducible, scriptable policies. Neither category fully addresses the need for a transparent and enforceable interpretation layer at the point of data ingestion, leaving users exposed to multiple failure modes summarized in Box 1, particularly in automated and batch-processing scenarios (Failure mode 6)[4].

OMIO is designed to fill this conceptual gap. Rather than introducing a new reader backend or visualization environment, OMIO acts as a policy-driven orchestration layer on top of existing readers. It defines explicit rules for axis semantics, metadata normalization, fallback behavior, and workflow-level operations, and enforces these rules uniformly across formats and acquisition systems.

In contrast to application-centric workflows, OMIO targets automated and programmatic analysis pipelines, where reproducibility, inspectability, and long-term robustness are critical. By separating semantic policy from format access and visualization, OMIO complements rather than replaces existing tools, providing a stable foundation for quantitative microscopy analysis across diverse software ecosystems.

The persistence of these issues despite the availability of standardized data models underscores the distinction between formal specification and effective semantic enforcement in practice, a gap that has been recognized for over a decade[1, 5, 18, 2].

## Design goals

OMIO is designed as a policy-driven infrastructure layer for microscopy image input and output. Its design is guided by a small set of explicit and enforceable goals that directly address recurring failure modes in quantitative microscopy workflows summarized in Box 1. These goals define the scope of OMIO and constrain its implementation choices.

### Canonical axis semantics

OMIO enforces a single canonical internal axis convention, TZCYX, across all supported input formats. Reader-specific axis orders are never propagated directly into downstream workflows. Instead, all image arrays are normalized at the API boundary such that time, depth, channel, and spatial dimensions are represented explicitly and in a fixed order.

A central aspect of this policy is that missing axes are inserted as singleton dimensions rather than being silently omitted, and existing singleton dimensions are preserved rather than implicitly squeezed away. This ensures that datasets with superficially different raw dimensionality remain structurally comparable once ingested by OMIO.

By replacing backend-specific axis conventions with a uniform and explicit representation, OMIO eliminates a major source of silent interpretation errors in downstream analysis code. This design goal directly addresses axis ambiguity, silent reordering, and shape-dependent behavior caused by implicit dimension handling (Failure modes 1 and 2 in Box 1).

### Metadata integrity and explicit fallbacks

OMIO prioritizes metadata integrity over convenience. Measurement-relevant metadata fields, in particular physical voxel sizes and units, are treated as first-class objects and normalized into a standardized representation. When metadata are incomplete, inconsistent, or ambiguous, OMIO applies controlled fallback strategies that are explicitly documented and communicated to the user via warnings.

OMIO never silently invents or infers physical metadata values. When a fallback value is applied, this decision is governed by an explicit policy and surfaced to the user through warnings, so that downstream quantitative interpretation remains auditable. All assumptions made during metadata normalization are surfaced, enabling users to assess the validity of quantitative results derived from the data. For Thorlabs RAW data, OMIO uses native Experiment.xml metadata when available and valid, but can fall back to explicit user-provided YAML sidecars when XML metadata are missing, incomplete, or inconsistent. This makes manual metadata repair a documented and inspectable part of the workflow rather than an implicit workaround. This design goal directly targets the loss or corruption of physical pixel sizes that commonly compromises quantitative microscopy analyses (Failure mode 3 in Box 1). Auxiliary or non-standard metadata are preserved separately to avoid loss of provenance while preventing contamination of core measurement fields.

### Separation of reader backends and semantic policy

OMIO deliberately separates format-specific reading from semantic normalization. Existing reader libraries are treated as interchangeable backends responsible only for extracting raw pixel data and available metadata. The interpretation, normalization, and validation of this information are handled exclusively by OMIO’s policy layer.

This separation directly addresses the reliance on undocumented, format-specific metadata heuristics that may change across library versions and undermine long-term reproducibility (Failure mode 4 in Box 1). By centralizing semantic decisions in a single policy layer, OMIO ensures that metadata interpretation is consistent across formats and independent of backend-specific implementation details.

### Open source and community-driven extensibility

OMIO is developed as an open source project with an explicit focus on community-driven evolution. Openness in this context refers both to the availability of the full source code and to an architectural design that actively encourages external contributions, feature requests, and extensions.

Support for microscopy file formats in OMIO is intentionally modular. Format-specific reader backends are implemented as interchangeable components with a well-defined interface, allowing new readers or reader extensions to be added without modifying the semantic policy layer. This design enables the community to contribute support for additional file formats, vendor-specific variants, or metadata extraction improvements in a targeted and maintainable manner.

When a file format or format variant is not yet supported, OMIO explicitly encourages users to contribute representative example files or reader implementations via the public issue tracker and contribution workflow. Reader support can be extended incrementally, either by adding new backend modules or by enriching existing metadata extraction routines. This approach avoids speculative or heuristic-based format inference and grounds support decisions in concrete data provided by the community.

By combining a stable semantic core with extensible reader backends, OMIO is designed to evolve alongside the rapidly changing microscopy software ecosystem, mitigating longterm fragmentation across tools and formats (Failure modes 4 and 6 in Box 1).

### Memory-aware operation

OMIO is designed to support large volumetric and time-resolved microscopy datasets that exceed available system memory. To this end, OMIO provides optional Zarr-based[19] backends that enable lazy loading and on-disk caching of image data. Data transfer into Zarr stores is performed incrementally to minimize peak memory usage. Disk-backed caches can be validated and reused across sessions, and users can place cache stores in custom locations, for example on local storage when source files reside on a remote server. This design goal supports reproducible and scalable analysis workflows by enabling consistent data handling across datasets of varying size, reducing the need for format-specific or dataset-specific memory management logic in downstream pipelines.

### Scalable visualization and Napari integration

OMIO is designed to support fast, correct, and scalable visualization of multidimensional microscopy data, with a particular focus on integration with the Napari viewer[13, 14]. A central goal is to ensure that axis semantics, physical scales, and dimensionality are represented correctly during visualization, independent of file format or dataset size.

To achieve this, OMIO exposes image data together with explicit axis labels and physical voxel sizes that can be consumed directly by Napari without additional user intervention. This design directly mitigates errors arising from inconsistent axis interpretation and metadata loss during visualization workflows (Failure modes 1, 2, and 5 in Box 1).

For large datasets, OMIO supports memory-efficient visualization workflows by combining Zarr-based memory mapping with optional parallelized data preparation. When enabled, data can be materialized into Zarr stores that allow chunked access and lazy loading. Preparation steps for Napari visualization, such as axis squeezing or reformatting for viewer compatibility, can be parallelized using Dask[20, 21] to reduce latency for interactive inspection.

This design goal ensures that OMIO serves not only as a reliable I/O and conversion layer, but also as a practical bridge between raw microscopy data and exploratory visualization. By integrating scalable data access patterns directly into the I/O layer, OMIO enables responsive inspection of large volumetric and time-resolved datasets on commodity hardware.

### Workflow-level semantics

OMIO operates at the level of scientific workflows rather than isolated files. In addition to single-image I/O, OMIO supports folder-based stacks, multi-file OME-TIFF series, explicit merging of datasets along defined axes, and flexible batch conversion of project trees following BIDS-like organizational principles[22].

By encoding these operations directly into the I/O layer, OMIO reduces the need for custom glue code and directly addresses workflow fragmentation and batch conversion inconsistencies that commonly arise in heterogeneous projects (Failure modes 5 and 6 in Box 1).

### Controlled OME-TIFF output

OMIO treats OME-TIFF writing as a controlled and semantically constrained operation. All output files adhere to the canonical axis ordering and include normalized metadata serialized into OME-XML. Decisions such as BigTIFF usage, compression, and filename provenance are handled deterministically based on data characteristics and available metadata.

This design goal ensures that OMIO outputs are immediately interoperable with standard microscopy tools and suitable as stable inputs for downstream analysis pipelines, preventing the reintroduction of semantic inconsistencies at the point of data export (Failure modes 1, 3, and 5 in Box 1).

### Implementation

OMIO is implemented as a modular Python package organized around a small public API and a larger set of internal helper routines. The implementation explicitly separates format access from semantic policy. Format specific readers are responsible only for extracting pixel data and any available metadata, while OMIO enforces axis semantics, metadata normalization, and workflow behavior uniformly across all backends.

### Architecture overview

At the highest level, OMIO exposes a small and deliberately constrained public API that defines the boundary between user-facing functionality and internal semantic enforcement. The public entry points are:

- imread: high-level reader for single files, lists of files, and folders, with optional Zarr materialization, folder stack handling, and merges.
- imwrite: controlled OME-TIFF writer that serializes OMIO-normalized metadata into OME-XML via tifffile.imwrite.
- imconvert: convenience wrapper implementing read plus write with a single call and consistent semantics.
- bids_batch_process: flexible batch processor for BIDS-like project trees[22] with explicit subject and folder-token matching rules.

Fig. 1 provides a schematic overview of this architecture, highlighting the strict separation between format-specific data extraction and centralized semantic normalization at the OMIO API boundary.

**Figure 1:**
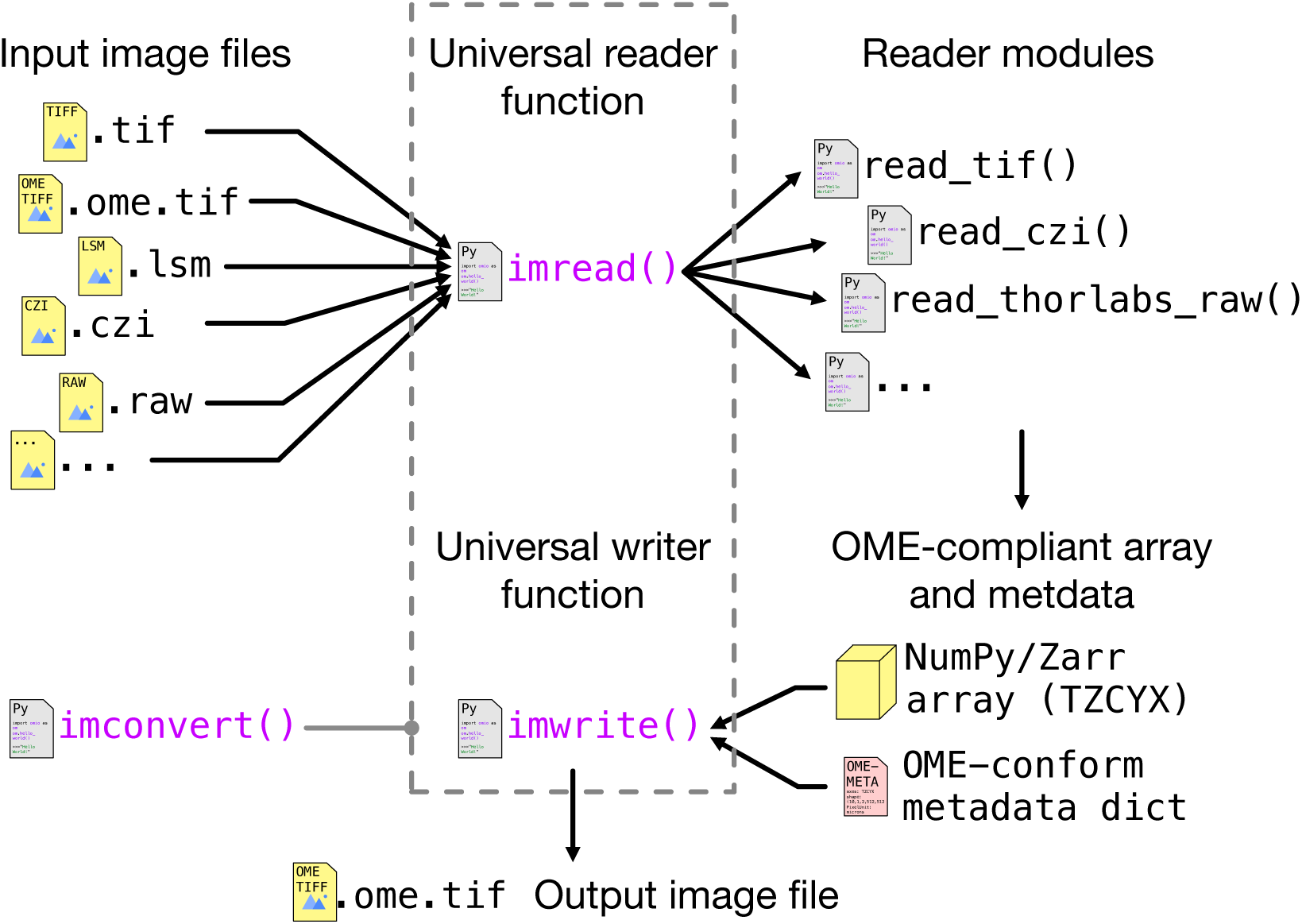
Architecture of the OMIO image I/O pipeline. Format-specific reader backends are responsible exclusively for low-level pixel and metadata extraction and are accessed through a single universal reader interface. All semantic decisions, including axis normalization, metadata standardization, and fallback handling, are enforced centrally at the OMIO API boundary. This separation yields a canonical OME-compatible in-memory representation with explicit axis semantics and validated metadata, which can be written deterministically to OME-TIFF without reintroducing backend-specific assumptions.

Internally, imread delegates file-level reading to a lightweight dispatch routine, which selects an appropriate format-specific backend based on explicit format cues such as file extensions and minimal structural inspection. These backends are responsible exclusively for low-level pixel and metadata extraction and do not perform any semantic interpretation or axis manipulation.

All format-specific readers share a common contract: they return an image object together with a standardized metadata dictionary. The image object is either a NumPy array[23] or a Zarr array[19], depending on whether Zarr materialization is requested. The associated metadata expose a normalized and explicitly structured key set, including an OME-compatible axis label string (axes), the corresponding array shape, measurement-relevant physical calibration fields such as PhysicalSizeX, PhysicalSizeY, and PhysicalSizeZ with associated units, as well as provenance information including the original file name, file type, and metadata source. Additional metadata that cannot be promoted unambiguously to measurement-relevant keys are preserved separately in a dedicated Annotations container.

The central invariant is that the final output of imread is normalized to a canonical OME-compatible axis order and is accompanied by metadata that are explicitly checked and corrected for internal consistency.

### Modular reader backends and community extensibility

OMIO is implemented as a modular I/O system in which format-specific functionality is isolated into dedicated reader backends with a strictly defined interface. Each backend is responsible exclusively for low-level data access, that is, extracting pixel arrays and exposing any available native metadata. Semantic interpretation, axis normalization, metadata validation, and workflow-level behavior are intentionally excluded from backend responsibilities and are handled centrally by OMIO’s policy layer. This separation between backend-specific extraction and centralized semantic enforcement is illustrated in Fig. 1. At the implementation level, format-specific backends are orchestrated through a lightweight dispatch mechanism within imread. Based on explicit and inspectable cues such as file extensions and minimal structural properties, OMIO selects an appropriate backend and delegates raw data extraction without applying any semantic transformations at this stage. Backend functions are required to return exactly two objects: an image container, represented either as a NumPy array or a Zarr array depending on configuration, and a raw metadata dictionary reflecting the information available in the source file. No backend is permitted to reorder axes, insert or remove dimensions, or otherwise modify semantic meaning, ensuring that all downstream transformations are applied uniformly and remain auditable.

This dispatch mechanism extends naturally from single-file inputs to collections of files and directory-based datasets. OMIO supports reading individual files, explicit lists of files, and folder-based inputs. When folders are provided, OMIO applies explicit and parameterized interpretation rules to determine whether the folder represents a collection of independent images, an ordered stack along a specified axis, or a collection of such stacks to be merged into higher-dimensional datasets. Crucially, these interpretations are never inferred implicitly from file naming conventions or directory layout alone, but are controlled by explicit user intent encoded at the API boundary.

A key semantic invariant enforced at this stage is that heterogeneous file layouts are never implicitly flattened. For example, paginated or multi-series image files are returned as structured collections of stacks rather than being concatenated into a single array by default. The return type of imread therefore reflects the logical structure of the input: either a single normalized image–metadata pair or a structured collection thereof. This design prevents silent loss of layout information and ensures that downstream processing can reason explicitly about dataset structure.

This modular dispatch architecture is a deliberate design choice to support extensibility and community-driven development. New file formats or format variants can be integrated by adding self-contained backend modules without modifying the core normalization pipeline. Similarly, vendor-specific metadata conventions can be accommodated by extending localized metadata extraction or standardization routines, while preserving the global semantic invariants enforced by OMIO.

Unsupported formats are handled conservatively. OMIO avoids heuristic inference or silent fallback behavior when encountering unknown layouts. Instead, the preferred extension pathway is the provision of representative example files or targeted backend contributions. In practice, adding support for a new format typically consists of implementing a small reader backend and, if necessary, augmenting metadata standardization helpers, without altering axis semantics, merge logic, or output guarantees.

By enforcing a strict boundary between backend-specific extraction, structured dispatch, and centralized semantic enforcement, OMIO’s implementation remains maintainable as format coverage grows. The open-source and community-oriented character of the project is thus embedded directly into the code structure itself, enabling external contributions while preserving internal consistency and long-term stability.

### Reading and axis normalization

All readers converge on a shared axis normalization pipeline. Conceptually, OMIO implements a deterministic mapping

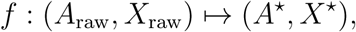

where *X*_raw_ is the raw array, *A*_raw_ its associated axis string, and *A*\* = TZCYX the canonical target axis order. Operationally, this mapping consists of two steps:

1. **Permutation** of existing axes into canonical order.
2. **Explicit insertion of missing axes** as singleton dimensions of length one.

Formally, let *π* be the permutation that reorders the axes present in *A*_raw_ into the order induced by *A*\*. OMIO applies this permutation to the array dimensions, followed by insertion of singleton dimensions for axes in *A*\* that are absent in *A*_raw_. The result is an array *X*\* whose rank equals *|A*\**|* and whose axis semantics are explicit and uniform across datasets.

This design avoids implicit squeezing and guarantees that downstream code can rely on a fixed dimensional interface, even when certain axes have length one. This deterministic axis normalization pipeline directly addresses axis ambiguity and silent reordering across reader backends, as well as shape-dependent behavior caused by implicit dimension squeezing (Failure modes 1 and 2 in Box 1).

When axes are missing or underspecified, OMIO inserts singleton dimensions instead of implicitly squeezing arrays. This behavior is enforced by helper routines that ensure consistency between axis labels and array shape. The guiding principle is that implicit assumptions are replaced by explicit representation. Arrays that differ only by the absence of a time, depth, or channel dimension are made structurally comparable by introducing explicit singleton axes, eliminating shape dependent behavior in downstream analysis.

This behavior can be stated formally by considering the intermediate result of the mapping *f* defined above. After permutation of existing axes into canonical order, let *A*_raw_ *⊆ A*\* denote the set of axis symbols present in the input (unchanged by permutation), and let *X*_perm_ be the resulting array. For each axis symbol *a ∈ A*\* *\ A*_raw_, OMIO inserts an explicit singleton dimension of length one at the position prescribed by *A*\*. Formally, if *X*_perm_ has shape (*s*_a1_*, . . ., s*_a_*_k_*) for the ordered axes *a*_1_*, . . ., a*_k_ *∈ A*_raw_, the final normalized array *X*\* is constructed such that

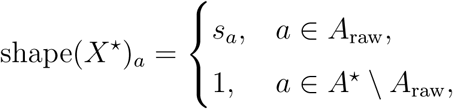

with axes ordered according to *A*\* = TZCYX. This guarantees that rank(*X*\*) = *|A*\**|* for all inputs, independent of the dimensional completeness of the original file.

By construction, no axis is ever removed or implicitly collapsed. Dimensions of length one remain explicit, and arrays that differ only by the absence of time, depth, or channel dimensions in the source representation are mapped to structurally comparable tensors. This eliminates shape-dependent branching in downstream analysis code and replaces implicit assumptions about dimensionality with an explicit and uniform representation. Together, these guarantees directly address axis ambiguity, silent reordering, and shape-dependent behavior caused by implicit dimension handling (Failure modes 1 and 2 in Box 1).

### Metadata standardization and integrity checks

OMIO enforces explicit metadata standardization as a central component of its semantic policy. The goal is not to maximize metadata extraction, but to ensure that a well-defined and measurement-relevant subset of metadata fields is represented consistently and remains quantitatively meaningful across formats and workflows. This policy-level separation between standardized and auxiliary metadata is illustrated schematically in Fig. 2.

**Figure 2:**
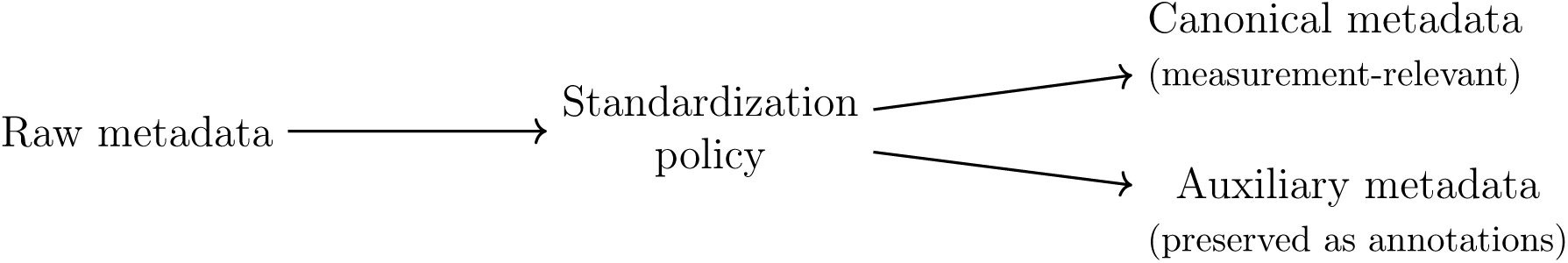
Metadata handling policy in OMIO. Raw metadata extracted from input files are processed by a standardization policy, which splits them into a canonical, measurement-relevant subset and auxiliary metadata preserved verbatim as annotations.

As shown in Fig. 2, raw metadata extracted from input files are never interpreted directly. Instead, OMIO applies a centralized standardization policy that promotes only a restricted, measurement-relevant subset to canonical keys, while all remaining information is preserved separately as auxiliary annotations.

OMIO’s metadata handling is governed by two complementary principles. First, a small set of measurement-relevant fields is promoted to stable, standardized keys with explicit semantics. Second, all remaining metadata are preserved without being allowed to contaminate or destabilize the standardized representation.

Measurement-relevant metadata include axis labels, array shape, physical voxel sizes, temporal sampling information where available, and essential provenance descriptors. These fields are normalized into a consistent schema that is independent of file format, vendor conventions, or acquisition software. All semantic interpretation and correction of these fields is applied centrally, after raw metadata extraction and axis normalization, ensuring that backend-specific quirks do not propagate into downstream analysis.

All additional metadata that cannot be mapped unambiguously onto this standardized schema are retained verbatim and stored separately as auxiliary annotations. This separation ensures that contextual or vendor-specific information is preserved for traceability and inspection, while preventing accidental use of non-standardized metadata in quantitative analysis. Where applicable, auxiliary metadata are structured in a form compatible with OME MapAnnotations to maintain interoperability with existing ecosystems.

After standardization, OMIO applies an explicit metadata integrity check that enforces internal consistency between axis semantics, array shape, and physical calibration. This check is applied uniformly to single images as well as to collections of images returned by layout-specific readers. Inconsistencies are corrected where possible, and otherwise surfaced to the user through warnings rather than being resolved implicitly.

A particularly high-risk metadata category is physical voxel size. Quantitative microscopy analysis critically depends on correct spatial calibration, yet voxel size information is frequently missing, incomplete, or inconsistently encoded across formats. OMIO therefore never silently infers voxel sizes. If reliable voxel size information cannot be extracted from the input metadata, OMIO requires either explicit user-provided values or applies documented default values, emitting warnings to make all assumptions visible and auditable.

By enforcing explicit metadata standardization, strict separation of auxiliary information, and transparent fallback behavior for critical quantitative fields, OMIO directly addresses the loss or corruption of measurement-relevant metadata that commonly occurs in heterogeneous microscopy workflows (Failure mode 3 in Box 1).

### Memory-aware backends and caching

OMIO supports returning Zarr arrays instead of NumPy arrays by setting zarr_store. Two main modes are implemented:

- in-memory Zarr stores for lightweight lazy workflows
- on-disk Zarr stores for large files and interactive exploration

To support interactive viewing and large dataset processing, OMIO implements incremental copying into Zarr stores. The primary low-level routine _copy_to_zarr_in_xy_ slices copies data one *XY* plane at a time, limiting peak RAM usage. On-disk Zarr stores contain OMIO metadata and cache manifests that allow existing stores to be validated and reused instead of rebuilt. The cache location can also be configured explicitly via zarr_store_path, which supports workflows in which large source files remain on network or external storage while the working cache is kept on a local disk.

For visualization, OMIO includes helper routines that derive inspection-oriented views from the canonical representation, including controlled removal of trivial singleton dimensions and an optional on-disk cache in a .omio_cache folder. These views enable interactive exploration of large datasets without modifying the normalized representation used for quantitative analysis. Zarr-backed inputs can also be written to OME-TIFF plane by plane, avoiding full-store materialization during export while preserving the controlled OME-TIFF output policy.

### Napari-compatible visualization pathways

Interactive visualization is treated in OMIO not as an auxiliary convenience, but as a first-class constraint that informs the design of the internal data representation. In particular, OMIO targets direct compatibility with the Napari viewer, which imposes strict requirements on array layout, axis semantics, physical scaling, and memory access patterns.

To satisfy these requirements without introducing viewer-specific logic into the analysis pipeline, OMIO enforces all semantic guarantees at the I/O boundary. Arrays returned by OMIO are normalized to a canonical OME-compatible axis order and are accompanied by standardized physical voxel sizes. As a result, image data can be passed directly to Napari with correct dimensional interpretation and spatial calibration, without post hoc reordering or manual adjustment.

For large volumetric and time-resolved datasets, OMIO supports Napari-compatible lazy access via optional Zarr-based backends. In this configuration, data are accessed at the slice level, such that only the currently viewed planes are loaded into memory. Zarr materialization is performed incrementally using slice-wise copy strategies to limit peak memory usage, enabling interactive inspection of datasets that exceed available RAM.

In addition to the canonical representation used for quantitative analysis, OMIO provides explicit visualization-oriented views that adapt the normalized data to Napari’s expected dimensionality. These views remove trivial singleton dimensions while preserving axis order and physical calibration. Importantly, this adaptation is performed deterministically and cached separately, ensuring that visualization convenience does not alter or obscure the canonical representation.

By treating Napari compatibility as an explicit architectural constraint rather than an external integration step, OMIO links robust semantic normalization with practical exploratory visualization. Viewer-specific requirements are satisfied without weakening the core guarantees of axis consistency, metadata integrity, or reproducibility enforced throughout the I/O pipeline.

### Writing OME-TIFF output

OMIO provides a controlled and policy-driven writing stage via imwrite, which serializes normalized image data and metadata into OME-TIFF files. Writing is treated not as a passive export step, but as the final enforcement point of OMIO’s semantic guarantees.

Before serialization, OMIO resolves output naming and file placement deterministically based on provenance information when available, ensuring reproducible file organization and avoiding silent overwriting. The writer explicitly determines whether the BigTIFF variant is required, based on an internal size estimate, so that large datasets are handled robustly without user intervention.

Crucially, imwrite never operates on unnormalized data. Arrays are first converted into the axis order required by the OME-TIFF specification, and all metadata written to disk are derived exclusively from OMIO’s standardized internal representation. For Zarr-backed inputs, this conversion can be performed as a plane-wise streaming operation so that the output remains a regular OME-TIFF file without requiring the full Zarr store to be materialized as a dense NumPy array. This guarantees that no backend-specific axis conventions, implicit dimension handling, or unvalidated metadata re-enter the data during export.

Measurement-relevant metadata such as PhysicalSizeX, PhysicalSizeY, PhysicalSizeZ, and TimeIncrement are written explicitly into the OME-XML payload with consistent unit handling. Auxiliary metadata that were preserved during reading and standardization are serialized separately as OME MapAnnotations, ensuring that contextual and provenance information remains accessible without contaminating the measurement-relevant metadata schema.

By enforcing canonical axes and validated metadata at write time, imwrite ensures that OME-TIFF files produced by OMIO are not merely syntactically valid, but semantically stable and analysis-ready. This prevents downstream tools from reintroducing ambiguity through implicit assumptions and closes the normalization loop established at read time.

### Workflow-level operations

OMIO implements workflow primitives that are common in practice but frequently reimplemented as lab specific scripts. These scripts typically embed implicit assumptions about directory layout, file naming, or acquisition conventions, and often mix format handling, semantic interpretation, and batch logic in a single, non-reusable implementation. As a result, equivalent workflows are repeatedly re-created across laboratories with subtle but consequential differences in behavior and semantics. OMIO instead lifts these operations to the level of dataset collections and project structure rather than individual files, enforcing a uniform semantic policy across heterogeneous inputs.

#### Folder reading and folder stacks

In addition to reading single files, imread supports folder inputs and can interpret them as:

- collections of independent image files
- folder stacks where files represent ordered slices along a specified axis
- collections of folder stacks that can be merged into higher dimensional arrays

The key behavior is that OMIO makes such interpretations explicit through parameters rather than relying on implicit filename heuristics.

#### Concatenation and merges along OME axes

OMIO supports explicit concatenation of multiple arrays along a selected OME axis via _merge_concat_along_axis. Given input arrays *X*_i_ with shapes *S*_i_, the routine constructs a target shape

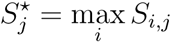

for all non merge axes *j*, and embeds each *X*_i_ into the corresponding slice of the output array. If zero padding is enabled, missing regions are filled deterministically with zeros.

This defines a well specified merge operator that avoids silent shape truncation and ensures reproducible behavior even when input arrays differ slightly in extent.

#### BIDS-like batch processing

`bids_batch_process` implements a deterministic traversal of project trees organized in a BIDS-like manner[22], with explicit matching rules for subject folders and flexible folder-token levels. In its default configuration, the function acts as an OMIO batch converter: as illustrated in Fig. 3, each discovered file is passed through OMIO’s unified read–normalize–write pipeline and written as an OME-TIFF output with standardized metadata and preserved provenance. Rather than relying on a fixed directory schema, the implementation uses explicit subject, folder-token, file-pattern, exclusion, and skip rules to identify eligible inputs. This enables reproducible conversion of large collections of microscopy datasets into a uniform representation suitable for downstream pipelines.

**Figure 3:**
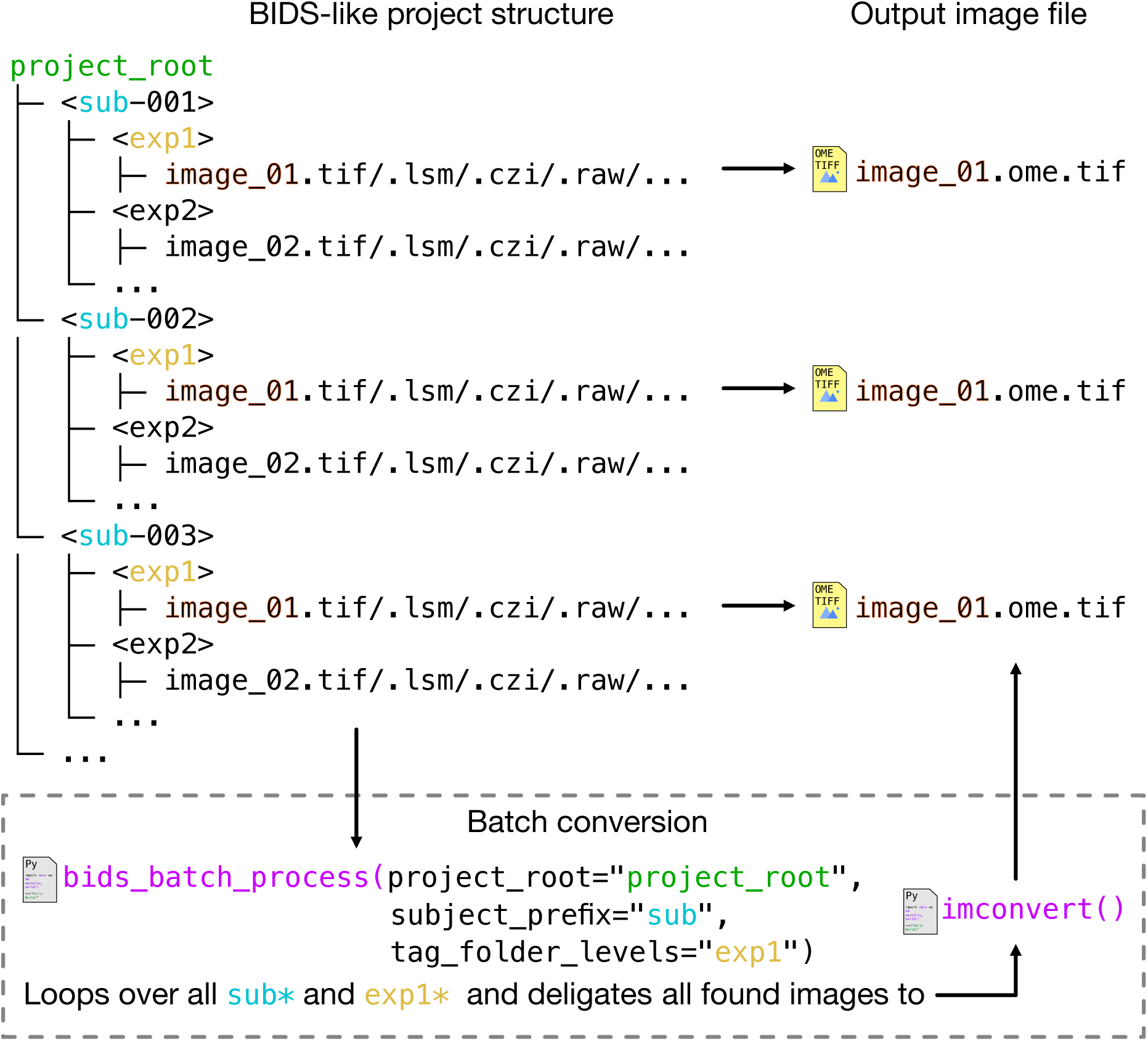
Deterministic BIDS-like batch processing workflow implemented by bids_batch_process. The schematic shows how OMIO performs project-level batch conversion of microscopy datasets organized in a BIDS-like directory structure. Input data are grouped by explicit subject and folder-token matching rules, while individual image files may exist in heterogeneous vendor-specific or standard formats (upper left). Batch processing is initiated by a single call to `bids_batch_process` (bottom center), which traverses the project tree using explicit matching rules and applies OMIO’s unified read–normalize–write pipeline by default. Optional processing callables can be inserted between reading and writing without changing the I/O policy. Each output image is written as an OME-TIFF file with canonical axis order (TZCYX) and normalized metadata (upper right). By operating on project structure rather than filenames or implicit heuristics, `bids_batch_process` enforces consistent semantic interpretation and enables reproducible, scalable processing of large microscopy collections.

`bids_batch_process(project_root=’data/raw’,`

`output_folder_name=’omio_processed’,`

`subject_prefix=’ID’,`

`tag_folder_levels=[(’exp1’, ’exp2’), (’confocal’,)])`

Beyond conversion, bids_batch_process can also operate as a lightweight batch processing orchestrator by accepting custom callables for processing steps between OMIO’s default loading and saving stages. This allows users to apply simple, explicit operations, such as projections, filtering, channel selection, or quality-control calculations, across entire project trees while retaining OMIO’s standardized I/O semantics and persistent run and error reports.

Taken together, these workflow-level primitives address fragmentation and inconsistency in batch processing pipelines by enforcing uniform semantic policies across datasets and processing steps, independent of input layout or acquisition provenance (Failure modes 5 and 6 in Box 1).

### Summary of enforcement points

The implementation contains three explicit enforcement points that are central for reproducibility:

- Axis normalization is applied after every read, before returning from the reader.
- Metadata standardization and schema checkup are applied post hoc, after axis operations.
- OME-TIFF writing always serializes canonical axes and normalized metadata and never writes unnormalized arrays.

These enforcement points ensure that semantic policy is applied consistently across formats and workflows.

### Evaluation

The evaluation is structured around the semantic risks summarized as failure modes in Box 1 and quantifies whether OMIO enforces the intended invariants across heterogeneous inputs and repeated conversion. It focuses on semantic correctness, robustness, and practical usability. Semantic validation includes round-trip tests in which data are read, written to OME-TIFF, and re-read, verifying invariants such as axis order, shape consistency, and physical pixel sizes. Robustness is assessed using a suite of heterogeneous inputs, including single-file TIFFs, multi-file OME series, paginated TIFFs, vendor-specific formats, and datasets with incomplete metadata. Memory behavior is evaluated by comparing NumPy-based and Zarr-based workflows on large volumetric time series, demonstrating the feasibility of interactive processing under constrained memory conditions. All tests were conducted using OMIO version v0.3.0[24] on a MacBook Pro 14-inch with an Apple M1 Max chip and 64 GB RAM running macOS Tahoe 26.5.

### Axis semantics and normalization behavior

To evaluate the correctness and robustness of OMIO’s axis handling, we assessed how reliably heterogeneous microscopy file formats are mapped onto a unified and explicit axis representation. The focus of this evaluation is semantic correctness and transparency of axis normalization, rather than raw I/O throughput. The core goal is to determine whether OMIO produces a consistent, analysis ready representation while making all assumptions about dimensionality explicit.

A heterogeneous set of microscopy image files was assembled, covering common formats with substantially different conventions regarding dimensional completeness, axis ordering, and metadata explicitness. The evaluation included generic TIFF, OME-TIFF, Zeiss LSM, and Zeiss CZI. These formats differ in whether all spatial and non spatial axes are explicitly encoded, in the ordering of dimensions as stored on disk, and in the extent to which axis semantics are described in machine readable metadata. Some files encode only spatial dimensions and implicitly assume missing axes, whereas others include time, depth, channel, or acquisition specific axes with conventions that are explicit in the vendor format but not necessarily aligned with analysis oriented axis order. The supplementary materials provide a detailed overview of the tested datasets.

Each file was processed as follows. First, a best effort estimate of the native axis string and raw array shape was obtained using format specific metadata inspection. For TIFF derived formats, including TIFF, OME-TIFF, and LSM, this step relied on *tifffile*[6] series axes when available, falling back to ImageJ metadata or OME XML dimension order when present, and otherwise using rank based synthesis for LSM when only the number of dimensions could be determined. For CZI files, raw axes were extracted using *czifile*[9]. Second, the file was read via OMIO’s imread, yielding a normalized NumPy array and standardized metadata including an explicit axis string. OMIO’s target representation for this evaluation was the fixed axis order

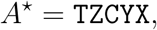

meaning that all datasets were expected to be represented in time, depth, channel, and spatial axes with missing dimensions made explicit as singleton axes of length one.

Three complementary metrics were computed to characterize OMIO’s axis normalization behavior in a way that matches the evaluation code and the plotted summary in Fig. 4. The first metric is an axis normalization success indicator per file *i*:

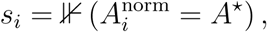

where 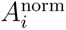 is the axis string returned by OMIO after normalization. The format wise success rate reported in Fig. 4a is the empirical mean

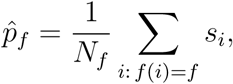

where *f* (*i*) denotes the format label of file *i* and *N*_f_ the number of files of that format.

**Figure 4:**
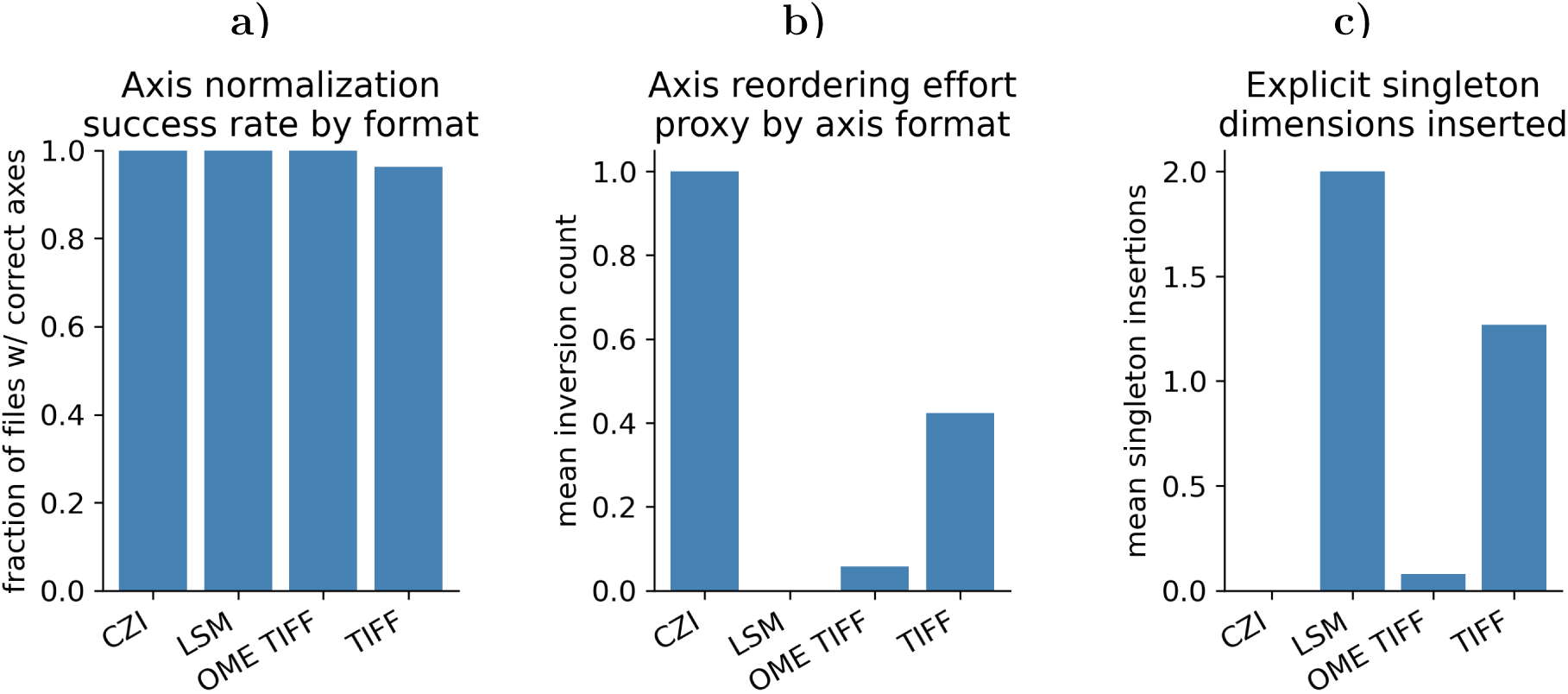
Axis normalization behavior across microscopy file formats. Shown is the behavior of OMIO’s axis normalization across native microscopy file formats, summarizing how reliably heterogeneous axis conventions are mapped to a unified, explicit target representation (TZCYX) and quantifying the normalization effort required for different formats. **Panel a)** Fraction of files per format whose normalized axis string exactly matches the target axis order. OMIO achieves consistent normalization for CZI, LSM, and OME-TIFF files, while generic TIFF files show a slightly reduced success rate due to incomplete or ambiguous native metadata. **Panel b)** Mean axis reordering effort per format, quantified as the number of pairwise inversions required to transform the native axis order into the normalized order. Higher values indicate stronger deviations between native acquisition-oriented conventions and the standardized analysis-oriented representation. CZI files require the most substantial reordering, whereas OME-TIFF and LSM files require little to none. **Panel c)** Mean number of explicitly inserted singleton dimensions per format. This reflects how often OMIO makes implicit dimensionality explicit, for example missing time, depth, or channel axes. Singleton insertions are frequent for TIFF and LSM files, rare for OME-TIFF, and absent for CZI, consistent with differences in metadata explicitness across formats.

This metric captures whether OMIO produces the intended explicit axis semantics without ambiguity.

The second metric quantifies a proxy for axis reordering effort. Let 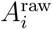 denote the best effort raw axis string and let 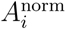 be the OMIO normalized axis string. The evaluation constructs the list of common axis symbols appearing in both strings, orders them as in the raw string, maps them to their positions in the normalized string, and computes the number of pairwise inversions in the resulting permutation. Formally, if this permutation vector is *P*_i_, the inversion count proxy is

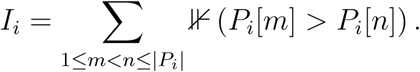

The plotted value in Fig. 4b is the mean inversion count per format,

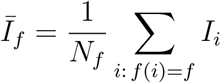

with undefined values excluded when raw axis information is unavailable. This metric does not measure computational cost, but it captures how strongly native axis conventions deviate from the standardized analysis oriented representation.

The third metric quantifies how often OMIO makes implicit dimensionality explicit by inserting singleton axes. For each file, the evaluation counts axis symbols that are present in 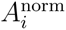 but absent in 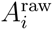, under the additional requirement that the corresponding normalized dimension length equals one. This metric should therefore be interpreted relative to the best-effort raw axis description, rather than as an assertion about the complete acquisition dimensionality encoded by the original experiment. Denoting this count by *S*_i_, Fig. 4c reports the per format mean

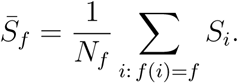

This metric captures how frequently OMIO must explicitly represent missing dimensions such as time, depth, or channel as singleton axes in order to ensure a uniform downstream interface.

Fig. 4 summarizes the results. In total, *N* = 270 files were evaluated, consisting of 241 OME-TIFF, 27 generic TIFF, one LSM, and one CZI file, with zero read errors in this set. The axis normalization success rate in Fig. 4a is 1.0 for OME-TIFF, LSM, and CZI, meaning all files in these groups were normalized to the target axis string. For generic TIFF, the success rate is 0.963, corresponding to 26 out of 27 files achieving the exact target axis string. This slight reduction is consistent with the fact that generic TIFF often lacks sufficient metadata to unambiguously infer non spatial dimensions, and it indicates conservative behavior rather than silent misinterpretation.

The axis reordering effort proxy in Fig. 4b highlights systematic differences between native conventions. CZI shows the largest mean inversion count in this set, with *Ī*_CZI_ = 1.0, reflecting acquisition oriented axis ordering that differs from the analysis oriented target representation. Generic TIFF exhibits moderate reordering effort, *Ī*_TIFF_ = 0.42, consistent with heterogeneous conventions across files. OME-TIFF is close to the target convention, with *Ī*_OME TIFF_ = 0.059, confirming near alignment between OMIO’s target representation and established OME standards. LSM shows negligible reordering effort, *Ī*_LSM_ = 0.0, consistent with the evaluation’s rank based canonicalization when explicit raw axis order is not available.

The singleton insertion metric in Fig. 4c further illustrates OMIO’s role in making hidden assumptions explicit. LSM requires *S̄*_LSM_ = 2.0 singleton insertions in this set, and generic TIFF requires *S̄*_TIFF_ = 1.269231, reflecting frequent absence of one or more non spatial dimensions in the native representation. OME-TIFF requires very few singleton insertions, *S̄*_OME TIFF_ = 0.078838, consistent with explicit axis metadata. CZI requires none in this set, *S̄*_CZI_ = 0.0, consistent with explicit encoding of dimensionality in the vendor format.

Taken together, this evaluation demonstrates that OMIO reliably normalizes axis semantics across heterogeneous microscopy file formats into a unified, explicit target representation. Differences in success rate, reordering proxy, and singleton insertion frequency reflect genuine differences in metadata completeness and native axis conventions across formats, rather than artifacts introduced by OMIO. By explicitly reordering axes when needed and introducing missing singleton dimensions only when justified by metadata absence, OMIO produces a uniform, analysis ready representation and reduces the risk of silent dimensionality errors in downstream pipelines.

### Round-trip invariance and metadata integrity

#### Voxel size integrity under round trip conversion

This evaluation isolates the behavior of physically meaningful voxel size metadata under a complete OMIO round trip, defined as a read–write–read sequence. The primary objective is to determine whether voxel sizes extracted and standardized by OMIO remain numerically invariant when written to OME-TIFF and subsequently re-read. A secondary objective is to characterize OMIO’s behavior in the presence of incomplete or internally inconsistent voxel size metadata, which are common in real microscopy datasets.

To test this under controlled and interpretable conditions, three synthetic single-file OME-TIFF datasets were generated. All datasets share identical image content, axis semantics, and array shape, but differ deliberately in their voxel size annotations. Each dataset uses axes TZCYX and an array shape (*T, Z, C, Y, X*) = (1, 5, 2, 32, 48) with random uint8 pixel values. The three voxel metadata conditions are as follows:

- **Complete metadata**: OME fields explicitly specify PhysicalSizeX = PhysicalSizeY = 0.19 *µ*m and PhysicalSizeZ = 2.0 *µ*m, with consistent units (µm). The TIFF resolution tag encodes the same lateral scale.
- **Partially missing metadata**: OME fields specify PhysicalSizeX = PhysicalSizeY = 0.19 *µ*m, while PhysicalSizeZ is intentionally omitted. The TIFF resolution tag again encodes 0.19 *µ*m laterally.
- **Conflicting metadata**: the TIFF resolution tag implies a lateral voxel size of 0.25 *µ*m, while the OME metadata explicitly specify PhysicalSizeX = PhysicalSizeY = 0.19 *µ*m and PhysicalSizeZ = 2.0 *µ*m. This mimics a common real-world conflict between multiple metadata sources.

Each dataset undergoes an identical processing pipeline. First, the file is read with OMIO using imread, yielding a NumPy array and standardized metadata. The returned metadata are then written back to disk using OMIO’s imwrite function in OME-TIFF format. Finally, the written file is read again with imread, and voxel sizes along the spatial axes are re-extracted.

For each spatial axis *a ∈ {X, Y, Z}*, the voxel size returned by OMIO on the initial read, 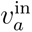, is compared to the voxel size obtained after re-reading the round-trip output, 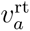. Absolute and relative errors are computed as

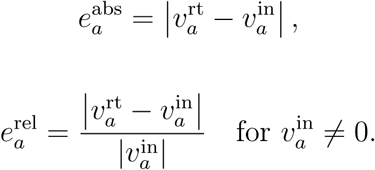

In addition, each axis is assigned a categorical status (preserved, changed, missing, introduced, or lost) based on the presence and numerical equality of voxel sizes before and after the round trip:

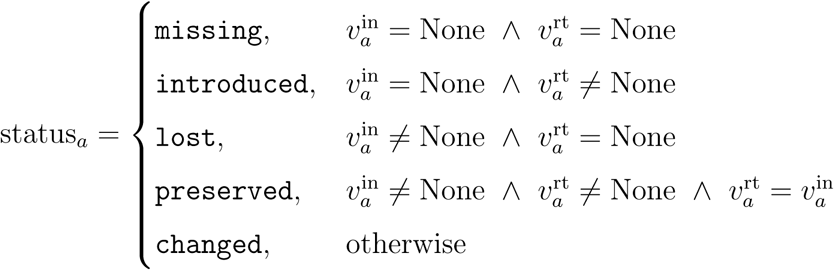

Dataset-level voxel completeness after the round trip is summarized by the indicator

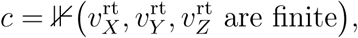

and the reported fraction of complete datasets is 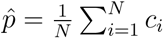, where *N* is the number of datasets.

The numerical results are reported in Table 2. Across all three datasets and all spatial axes, the maximum absolute and relative errors are zero, that is 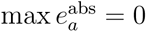 and 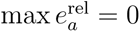 wherever a voxel size is defined. Concretely, the complete dataset preserves (0.19, 0.19, 2.0)*, µ*m exactly under the round trip. In the conflicting dataset, OMIO resolves the discrepancy deterministically in favor of the OME-encoded lateral voxel size 0.19*, µ*m and preserves this value exactly after writing and re-reading, demonstrating stable conflict resolution within OMIO’s standardization logic.

**Table 2:** Round-trip invariants of voxel size metadata across OMIO conversion. Reported are voxel size metadata before and after a read–write–read round trip through OMIO for synthetic OME-TIFF test datasets with complete, partial, and conflicting metadata. For each dataset and spatial axis (X, Y, Z), the standardized voxel size after the initial OMIO read, the voxel size after round trip, and a categorical status are listed. Preserved entries indicate exact numerical agreement between the standardized input representation and the round-trip representation. In the partially missing condition, the absent axial spacing is replaced by OMIO’s explicit fallback value and is then preserved under round trip. The absence of non-zero deviations demonstrates that voxel sizes, once standardized by OMIO, behave as strict round-trip invariants.

| File | Axis | Voxel<br>in $\mu\text{m}$ | Voxel roundtrip<br>in $\mu\text{m}$ | abs./rel.<br>error | Status |
| --- | --- | --- | --- | --- | --- |
| Complete metadata | X | 0.19 | 0.19 | 0/0 | preserved |
|  | Y | 0.19 | 0.19 | 0/0 | preserved |
|  | Z | 2.0 | 2.0 | 0/0 | preserved |
| Partially missing metadata | X | 0.19 | 0.19 | 0/0 | preserved |
|  | Y | 0.19 | 0.19 | 0/0 | preserved |
|  | Z | 1.0 | 1.0 | 0/0 | preserved |
| Conflicting metadata | X | 0.19 | 0.19 | 0/0 | preserved |
|  | Y | 0.19 | 0.19 | 0/0 | preserved |
|  | Z | 2.0 | 2.0 | 0/0 | preserved |

In the partially missing dataset, OMIO returns an axial voxel size of 1.0*, µ*m already on the initial read, despite the absence of PhysicalSizeZ in the input metadata. In this condition, OMIO applies its explicit fallback policy for missing axial spacing and emits a warning, thereby making the assumption auditable rather than silent. This value is subsequently written and preserved under the round trip without numerical deviation, and the evaluation confirms that this standardized value is treated as a round-trip invariant. The voxel completeness summary shown in Table 2 reflects this behavior: all three datasets have finite *X*, *Y*, and *Z* voxel sizes after the round trip, yielding a completeness fraction *p*^ = 1.0.

Overall, this evaluation demonstrates that voxel sizes behave as strict numerical invariants once standardized by OMIO, even in the presence of conflicting or incomplete metadata. At the same time, it explicitly documents OMIO’s current policy of supplying a default axial spacing when *Z* metadata are missing. The test therefore characterizes both the stability of voxel sizes under conversion and the effective semantics of OMIO’s metadata standardization, providing a transparent basis for interpreting physical calibration in downstream quantitative analyses.

#### Global round-trip consistency and idempotence

Building on the voxel-specific analysis above, this evaluation tests whether OMIO’s structural image representation is idempotent under repeated conversion, meaning that the representation produced by imread remains unchanged after an OMIO write and re-read cycle with respect to axis semantics, array geometry, and pixel values. In addition, a selected subset of standardized metadata fields is evaluated separately through a hash-based consistency check.

The test operates on heterogeneous real microscopy inputs scanned from the example data root, spanning Zeiss CZI, Zeiss LSM, generic TIFF, and OME-TIFF. The evaluation focuses on integer data to avoid floating-point tolerance issues and to directly test lossless behavior. For each eligible dataset, the pipeline is:

1. Read input via imread to obtain (*I*^in^*, M* ^in^).
2. Write OME-TIFF via imwrite.
3. Read written file via imread to obtain (*I*^rt^*, M* ^rt^).

Four invariants are then evaluated:

- **Axis semantics**: *b*_axes_ = *K*(*M* ^in^[axes] = *M* ^rt^[axes])
- **Shape**: *b*_shape_ = *K*(shape(*I*^in^) = shape(*I*^rt^))
- **Pixel values**: using the maximum absolute difference

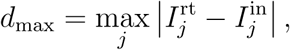

with *b*_pix_ = *K*(*d*_max_ = 0); *j* indexes voxels of the flattened arrays after confirming shape equality.

- **Standardized metadata stability**: a hash equality check on a fixed key subset,

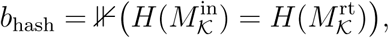

where *H* denotes a deterministic hash after canonical serialization of the selected metadata subset, and *K* includes axes, voxel sizes and units, and selected timing fields. The metadata hash check was used as an auxiliary consistency control and is not included in the plotted structural idempotence score.

A dataset is counted as structurally and pixel-wise idempotent under round trip if

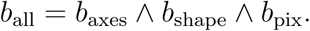

Fig. 5 summarizes the structural and pixel-level outcomes. Fig. 5a reports, for each input format, the fraction of evaluated datasets with *b*_all_ = 1. All shown formats reach a fraction of 1.0 in this run, indicating that every evaluated dataset preserved axis semantics, shape, and pixel values under the round trip. Fig. 5b complements this by showing the empirical distribution of sampled pixel differences Δ*I* = *I*^rt^ *— I*^in^. The histogram mass is entirely concentrated at Δ*I* = 0, consistent with *d*_max_ = 0 for the evaluated set. Together, these results show that OMIO’s conversion to OME-TIFF is lossless with respect to integer pixel data and preserves the structural semantics that OMIO standardizes, supporting the claim that OMIO does not introduce accumulating transformations under repeated use.

**Figure 5:**
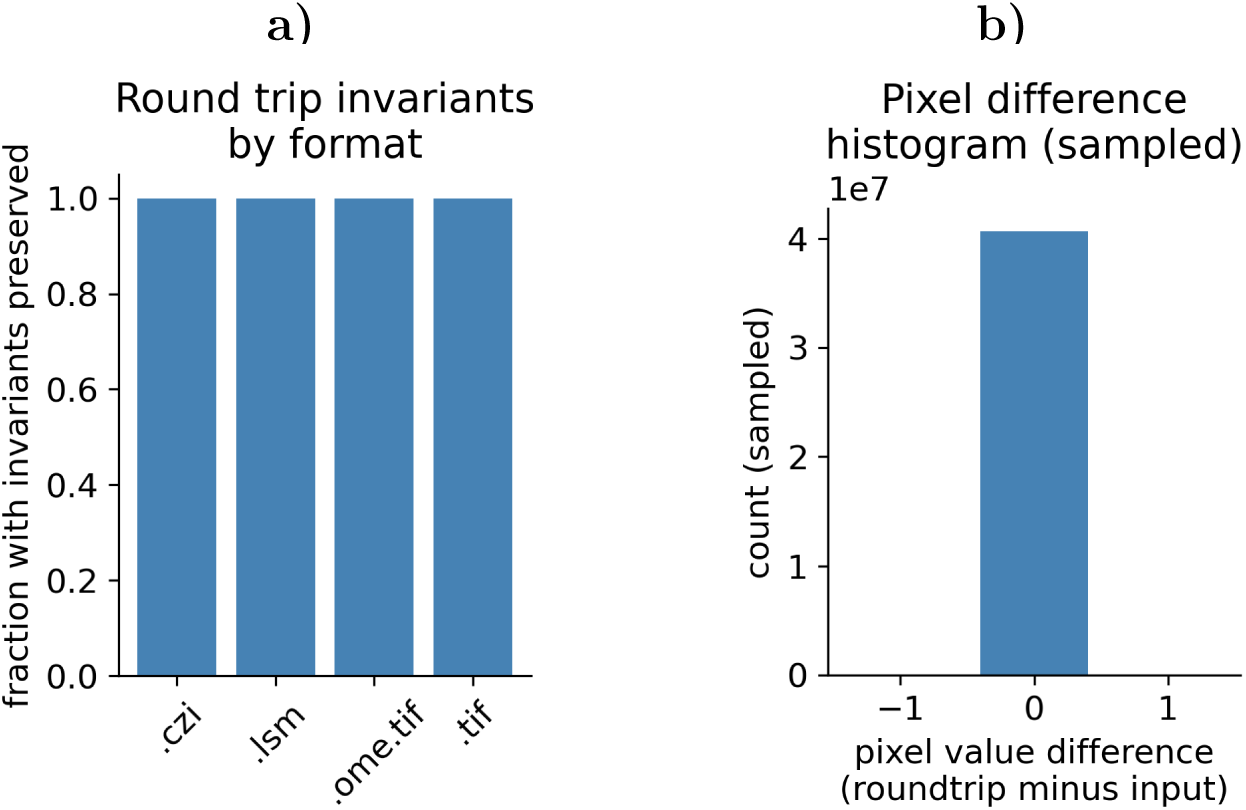
Round-trip invariants across native microscopy formats. Shown is the behavior of OMIO under a read–write–read round trip through OME-TIFF for heterogeneous microscopy file formats, evaluating whether axis semantics, array shape, and pixel values are preserved. **Panel a)** Fraction of evaluated datasets per input format for which OMIO preserves axis semantics, array shape, and pixel values under a read–write–read cycle. For each format (CZI, LSM, OME-TIFF, TIFF), the fraction equals 1.0 in this run, indicating that all tested datasets satisfy *b*_all_ = 1. **Panel b)** Histogram of sampled pixel value differences Δ*I* = *I*^rt^ *— I*^in^ for the same round trip, showing all sampled differences at Δ*I* = 0, consistent with zero maximum absolute pixel error. Source: OMIO evaluation, this work.

### Memory behavior and scalability

To evaluate the practical scalability of OMIO and its suitability for large volumetric data, we assessed both latency and memory behavior during image loading under different access strategies. The focus of this evaluation is not raw throughput in isolation, but the trade-off between time to first usable data and transient memory consumption, which is critical for interactive analysis and resource-constrained environments.

For this purpose, a set of synthetic single-file OME-TIFF datasets was generated with controlled size and content. All datasets follow the standardized axis order *TZCY X* and store unsigned 16-bit integer pixel values. The synthetic image content is generated deterministically using a pseudo-random number generator, drawing values uniformly from the full uint16 range [0, 65535]. This ensures non-trivial data while avoiding structure that could bias compression or caching behavior. Physical voxel sizes are encoded in the OME metadata with lateral pixel sizes of 0.19*, µ*m and an axial spacing of 2.0*, µ*m, reflecting typical two-photon microscopy acquisitions.

Three dataset scales were evaluated. The small dataset (S) has shape (*T, Z, C, Y, X*) = (1, 8, 1, 512, 512), corresponding to approximately 2.1 *×* 10^6^ voxels. The medium dataset (M) has shape (1, 16, 1, 768, 768), corresponding to approximately 9.4 *×* 10^6^ voxels.

The large dataset (L) has shape (1, 32, 1, 1024, 1024), corresponding to approximately 3.36 *×* 10^7^ voxels. Each dataset was written once to disk as a compressed OME-TIFF file and subsequently read repeatedly using three OMIO workflows: a NumPy-backed read (imread(…, zarr_store=None)), an in-memory Zarr representation (zarr_store= "memory"), and a disk-backed Zarr representation (zarr_store="disk").

Two primary metrics were measured. First, the time to first slice (TTFS), defined as the elapsed time until the first spatial plane becomes accessible. Formally, for a given workflow,

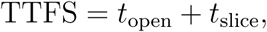

where *t*_open_ denotes the execution time of the imread call and *t*_slice_ the time required to access a single *Y X* plane at index (*t* = 0, *z* = 0, *c* = 0). This metric captures the latency experienced by an interactive user before meaningful data can be inspected.

Second, transient memory usage was quantified via the increase in resident set size (RSS) of the Python process during the TTFS measurement window. For each run, the RSS was recorded immediately before the operation and tracked continuously thereafter. The reported value is

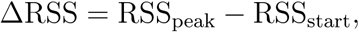

where RSS_peak_ is the maximum observed RSS during the operation and RSS_start_ the baseline RSS prior to loading. This differential metric isolates the memory impact attributable to the read and initial access itself. Measurements were repeated three times per condition and dataset size, with explicit garbage collection between repeats to reduce cross-run interference.

The top panel of Fig. 6 shows the resulting TTFS values. Across all workflows, TTFS increases monotonically with dataset size. The NumPy workflow exhibits the lowest latency at all scales, with TTFS of approximately 6–8 ms for S, *∼* 8 ms for M, and *∼* 23 ms for L. In contrast, the in-memory Zarr workflow shows higher TTFS, increasing from roughly 45 ms (S) to *∼* 90 ms (M) and *∼* 250 ms (L). The disk-backed Zarr workflow incurs the largest latency, with TTFS of approximately 55 ms (S), *∼* 145 ms (M), and *∼* 350 ms (L), reflecting filesystem access and chunk initialization overhead. These results illustrate the expected trade-off between eager and lazy access: NumPy minimizes latency by fully materializing data, whereas Zarr-based workflows defer data loading at the cost of increased initial access time.

**Figure 6:**
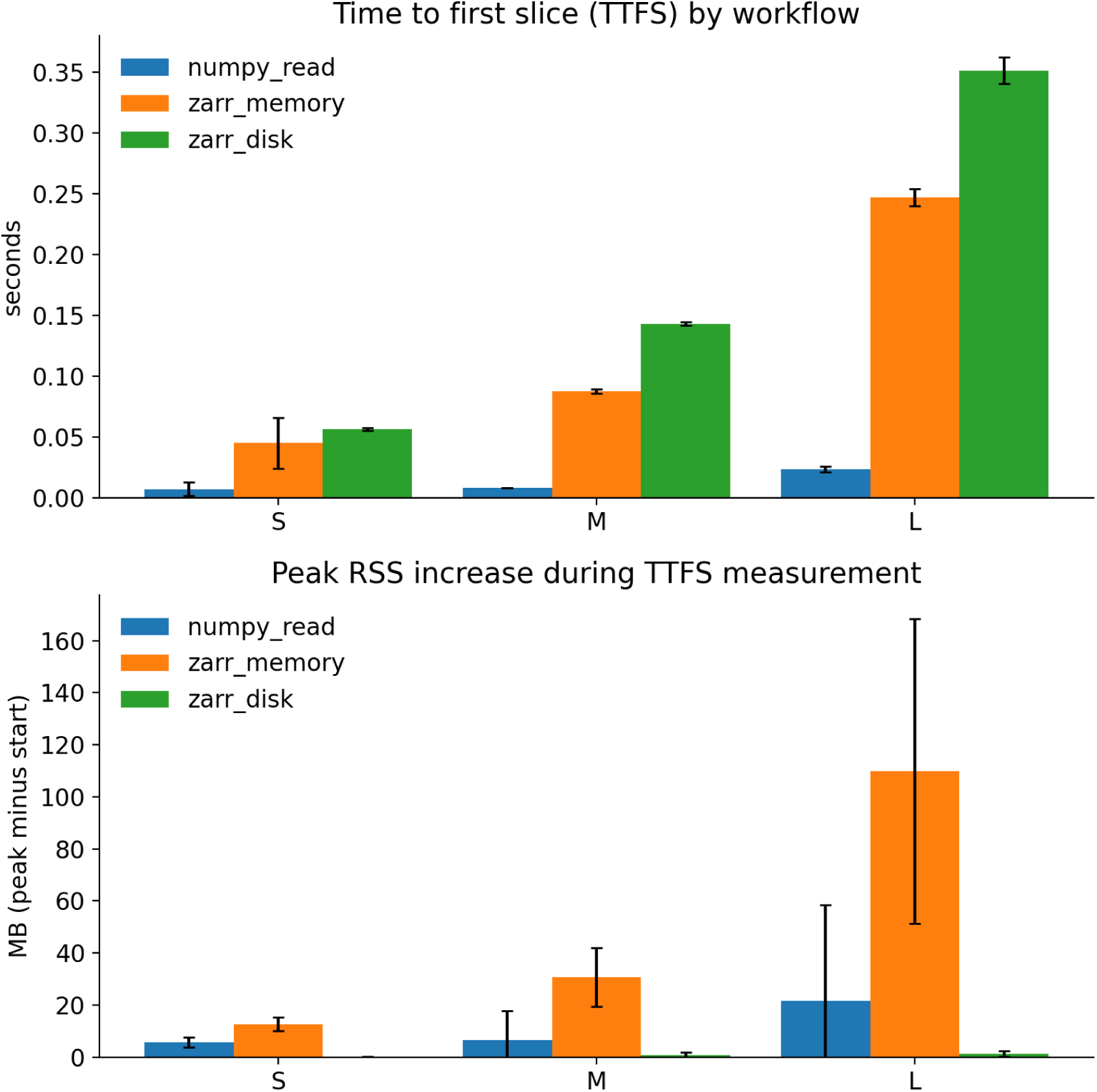
Memory behavior and scalability of OMIO across different access workflows. Compared are latency and transient memory usage when reading synthetic OME-TIFF datasets of increasing size using three OMIO workflows: full NumPy materialization (`zarr_store=None`), inmemory Zarr (`zarr_store="memory"`), and disk-backed Zarr (`zarr_store="disk"`). Top panel: time to first slice (TTFS), defined as the sum of image open time and access to the first *Y X* plane, shown for small (S), medium (M), and large (L) datasets. Bottom panel: peak increase in process resident set size (RSS) during the TTFS measurement, reported as ΔRSS = RSS_peak_ *—* RSS_start_. Error bars denote the standard deviation over repeated runs. Disk-backed Zarr shows near-zero RSS increase across all dataset sizes, reflecting chunked, lazy access with minimal memory retention, while NumPy and in-memory Zarr exhibit increasing memory usage with dataset size. Negative excursions due to measurement noise around zero are clipped at zero for visual clarity.

The bottom panel of Fig. 6 summarizes the corresponding transient memory behavior. For the NumPy workflow, ΔRSS increases with dataset size, from approximately 5 MB (S) to *∼* 7 MB (M) and *∼* 20 MB (L), consistent with increased memory pressure during dense array access. The measured RSS increase should be interpreted as a process-level transient memory metric rather than as a direct byte-level estimate of the stored array size, because it is affected by allocator behavior, caching, and memory reuse. The in-memory

Zarr workflow shows a substantially larger and more variable memory footprint, with mean ΔRSS of roughly 12 MB (S), *∼* 30 MB (M), and exceeding 100 MB for the large dataset, reflecting the additional overhead of chunked storage structures allocated in RAM alongside pixel data. In contrast, the disk-backed Zarr workflow exhibits ΔRSS values clustered near zero across all dataset sizes, with mean increases on the order of 0–2 MB. Small fluctuations around zero arise from measurement noise in RSS sampling rather than sustained memory allocation and are clipped at zero in the visualization for clarity.

Taken together, Fig. 6 demonstrates that OMIO provides predictable and tunable memory behavior. Users can trade increased TTFS for dramatically reduced memory pressure by selecting disk-backed Zarr access, enabling interactive inspection of datasets that would otherwise exceed available RAM. At the same time, NumPy-backed and inmemory Zarr workflows remain available for scenarios where minimal latency or repeated dense access outweigh memory constraints.

## Limitations

OMIO is intentionally conservative in situations where file formats or metadata do not provide sufficient information for unambiguous semantic normalization. In particular, OMIO avoids speculative inference of axis semantics or physical calibration. If essential metadata such as voxel sizes, axis ordering, or acquisition dimensionality are missing or internally inconsistent, OMIO either applies explicitly documented fallback values or requires user-provided input. While this behavior may require additional configuration effort, it ensures that quantitative interpretations remain auditable and that silent errors are avoided.

The degree of memory efficiency achievable during reading and conversion depends partly on the capabilities of the underlying reader backends. Some formats or third-party libraries require full materialization of image data in memory before conversion to a Zarr-backed representation is possible. In such cases, OMIO cannot fully eliminate peak memory usage, although it still provides a uniform downstream representation once data have been materialized.

OMIO does not aim to be a comprehensive replacement for specialized file format libraries or vendor-specific SDKs. Instead, it operates as an orchestration and normalization layer that builds on existing readers. Consequently, support for new or uncommon file formats depends on the availability of representative example data and backend implementations. Formats that are poorly documented or rely on proprietary encodings may require community contributions to enable robust support.

Finally, OMIO prioritizes semantic correctness and reproducibility over raw I/O throughput. In scenarios where maximal read performance is the primary objective and semantic normalization is unnecessary, lower-level libraries may provide faster access. OMIO is designed for workflows in which correctness, transparency, and interoperability outweigh minimal I/O latency.

## Availability and reproducibility

OMIO is an open-source Python package developed and maintained on GitHub and distributed via standard Python packaging mechanisms. The source code is publicly available at https://github.com/FabrizioMusacchio/omio, released under the GNU General Public License v3.0 (GPL-3.0), and can be installed directly from the Python Package Index using pip. All released versions of OMIO are versioned and archived with persistent identifiers to ensure long-term reproducibility. In addition, a curated example dataset is hosted on Zenodo (DOI: 10.5281/zenodo.18078231)[25], providing synthetic and representative microscopy files that can be used to test, evaluate, and reproduce the behavior described in this work. The example dataset is distributed under the Creative Commons Attribution 4.0 International license (CC BY 4.0).

Comprehensive user and developer documentation is provided via Read the Docs at https://omio.readthedocs.io/, including installation instructions, usage examples, API reference, executable tutorials, and contribution guidelines. The documentation and the GitHub repository both include explicit contribution guidelines and a code of conduct, outlining how users can report issues, request features, contribute new reader backends, or extend metadata handling in a structured and community-driven manner.

Together, the open availability of the source code, documented installation and usage workflows, archived example data, explicit licensing, and clear contribution pathways ensure that OMIO can be evaluated, reproduced, and extended by the broader microscopy and scientific Python communities.

### Prior use in lab workflows

Pre-release versions (prototype implementations predating the first public release) and early versions of OMIO were used in our laboratory in microscopy data conversion and analysis workflows supporting several projects and associated outputs, including published and preprint manuscripts as well as dataset releases [26, 27, 28, 29, 30, 31, 32, 33, 34]. This prior practical use informed OMIO’s explicit semantic policies and its focus on reproducible, auditable normalization at the I/O boundary.

## Summary

OMIO addresses a persistent and largely structural problem in quantitative microscopy workflows: the lack of a unified, explicit, and reproducible interface for image input and output across heterogeneous file formats and metadata conventions. Rather than introducing a new file format or replacing existing reader libraries, OMIO establishes a policy-driven I/O layer that enforces canonical axis semantics, standardized metadata handling, and controlled output behavior at the API boundary.

The evaluations presented in this work demonstrate that OMIO reliably normalizes axis semantics across diverse microscopy formats, preserves physically meaningful voxel size metadata under round-trip conversion, and offers predictable memory behavior for large datasets through optional Zarr-backed access. These results indicate that OMIO succeeds in making implicit assumptions explicit, reducing silent failure modes, and providing a stable, analysis-ready representation that downstream code can rely on without format-specific branching.

A central contribution of OMIO lies in its explicit separation of concerns. Format-specific readers are treated as interchangeable backends responsible only for data extraction, while semantic interpretation, normalization, and validation are enforced uniformly by OMIO itself. This architectural choice enables consistent behavior across formats, simplifies maintenance, and allows format coverage to expand without compromising semantic guarantees. At the same time, the modular design supports community-driven extension, encouraging contributions of new reader backends and metadata handling logic grounded in real-world example data rather than heuristic inference.

OMIO is not intended as a comprehensive replacement for specialized file format libraries or visualization tools. Instead, it occupies a deliberately narrow but critical layer between raw data acquisition and downstream analysis, providing controlled I/O semantics, reproducible workflow primitives, and interoperability with established tools such as OMETIFF-based pipelines and interactive viewers. In this role, OMIO complements existing ecosystems rather than competing with them.

Overall, OMIO demonstrates that enforcing explicit semantic policy at the I/O layer is both feasible and beneficial for microscopy workflows. By combining canonical axis semantics, metadata integrity, memory-aware operation, and workflow-level abstractions within an open and extensible framework, OMIO provides a practical foundation for reproducible and scalable image analysis in scientific Python environments.

## Acknowledgements

We thank the developers and maintainers of the open-source software ecosystem that OMIO builds upon, in particular NumPy[23], Zarr[19], tifffile[6], czifile[9], napari[14], and Dask[20, 21]. Their foundational work enables OMIO’s functionality and integration into the broader scientific Python landscape. We also acknowledge the creators of the openly licensed example datasets used during development and testing. Dataset-specific license information and required attributions are provided in the OMIO Zenodo example data record and accompanying README files[25].

MF received funding from the European Research Council (ERC; MicroSynCom 865618) and the German Research Foundation (DFG; SFB1089 C01, B06; SPP2395).

## Supplementary Information

**Table 3:**
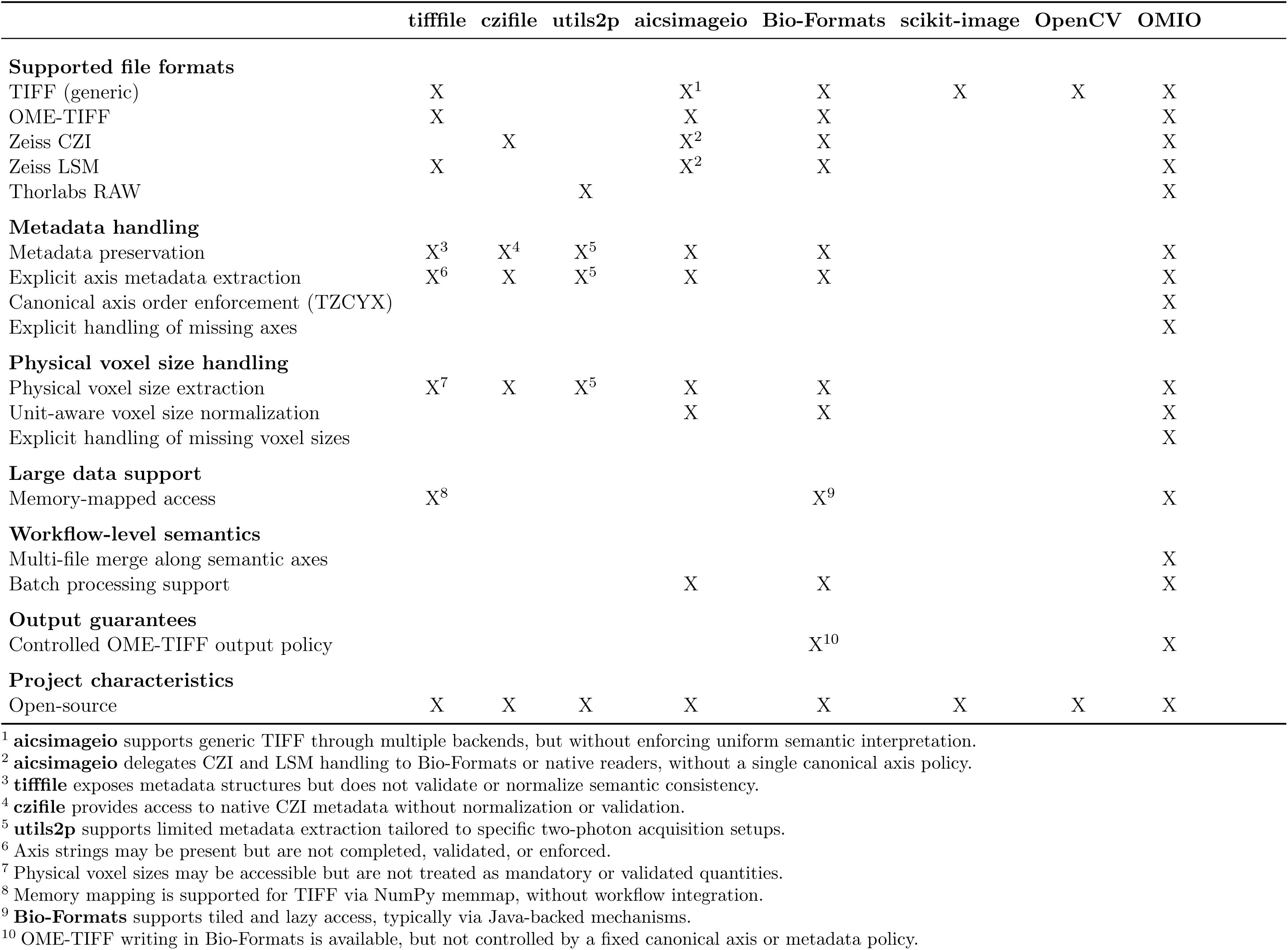
Extension of Table 1: Feature comparison of microscopy I/O libraries.

## References

[1] M. Linkert, C. T. Rueden, C. Allan, J.-M. Burel, W. Moore, A. Patterson, B. Loranger, J. Moore, C. Neves, D. MacDonald, A. Tarkowska, C. Sticco, E. Hill, M. Rossner, K. W. Eliceiri and J. R. Swedlow. ‘Metadata matters: access to image data in the real world’. In: Journal of Cell Biology 189.5 (May 2010), pp. 777–782. doi: 10.1083/jcb.201004104. eprint: https://rupress.org/jcb/article-pdf/189/5/777/1932292/jcb_201004104.pdf.

[2] J. R. Swedlow and K. W. Eliceiri. ‘Open source bioimage informatics for cell biology’. In: Trends in Cell Biology 19.11 (2009). Special Issue – Imaging Cell Biology, pp. 656– 660. doi: 10.1016/j.tcb.2009.08.007.

[3] I. G. Goldberg, C. Allan, J.-M. Burel, D. Creager, A. Falconi, H. Hochheiser, J. Johnston, J. Mellen, P. K. Sorger and J. R. Swedlow. ‘The Open Microscopy Environment (OME) Data Model and XML file: open tools for informatics and quantitative analysis in biological imaging’. In: Genome Biology 6.5 (2005), R47. doi: 10.1186/gb-2005-6-5-r47.

[4] M. Huisman, M. Hammer, A. Rigano, U. Boehm, J. J. Chambers, N. Gaudreault, A. J. North, J. A. Pimentel, D. Sudar, P. Bajcsy, C. M. Brown, A. D. Corbett, O. Faklaris, J. Lacoste, A. Laude, G. Nelson, R. Nitschke, D. Grunwald and C. Strambio-De-Castillia. A perspective on Microscopy Metadata: data provenance and quality control. 2021. arXiv: 1910.11370 [q-bio.QM].

[5] M. D. Wilkinson et al. ‘The FAIR Guiding Principles for scientific data management and stewardship’. In: Scientific Data 3.1 (2016), p. 160018. doi: 10.1038/sdata.2016.18.

[6] C. Gohlke. cgohlke/tifffile: v2025.12.20. Version v2025.12.20. 2025. doi: 10.5281/zenodo.18005696.

[7] CellProfiler developers. python-bioformats. Version 1.5.2. 2019.

[8] J. M. Brown, J. Sherman, toloudis, M. Swain-Bowden, B. Chaudhuri, G. Johnson, R. Casero and D. Hürlimann. AllenCellModeling/aicsimageio: Mosaics, Xarray, and Writers, Oh My! Version v4.0.0. 2021. doi: 10.5281/zenodo.4906609.

[9] C. Gohlke. czifile. Version v2019.7.2.1. 2019.

[10] NeLy-EPFL. utils2p. Version 1.0.2. 2024.

[11] J. Schindelin, I. Arganda-Carreras, E. Frise, V. Kaynig, M. Longair, T. Pietzsch, S. Preibisch, C. Rueden, S. Saalfeld, B. Schmid et al. ‘Fiji: an open-source platform for biological-image analysis’. In: Nature Methods 9.7 (2012), pp. 676–682. doi: 10.1038/nmeth.2019.

[12] C. T. Rueden, J. Schindelin, M. C. Hiner, B. E. DeZonia, A. E. Walter, E. T. Arena and K. W. Eliceiri. ‘ImageJ2: ImageJ for the next generation of scientific image data’. In: BMC Bioinformatics 18.1 (2017), p. 529. doi: 10.1186/s12859-017-1934-z.

[13] N. Sofroniew, et al. napari: a multi-dimensional image viewer for Python.

[14] N. Sofroniew, T. Lambert, G. Bokota, J. Nunez-Iglesias, P. Sobolewski, A. Sweet, L. Gaifas, K. Evans, A. Burt, D. Doncila Pop, K. Yamauchi, M. Weber Mendonça, J. Rodríguez-Guerra, L. Liu, G. Buckley, W.-M. Vierdag, A. Anderson, T. Monko, C. Willing and R. Zhao. *napari. a multi-dimensional image viewer for Python*. Version v0.7.0a0. 2025. doi: 10.5281/zenodo.18089604.

[15] Oxford Instruments. *Imaris*. Version 9.7. Commercial microscopy image analysis software. Accessed 2025.

[16] Carl Zeiss Microscopy GmbH. ZEISS ZEN Microscopy Software. https://www.zeiss.com/metrology/en/software/zeiss-zen-core.html. Accessed 2025.

[17] Leica Microsystems. *Leica LAS X*. Commercial microscopy software.

[18] R. A. Poldrack, C. I. Baker, J. Durnez, K. J. Gorgolewski, P. M. Matthews, M. R. Munafò, T. E. Nichols, J.-B. Poline, E. Vul and T. Yarkoni. ‘Scanning the horizon: towards transparent and reproducible neuroimaging research’. In: Nature Reviews Neuroscience 18.2 (2017), pp. 115–126. doi: 10.1038/nrn.2016.167.

[19] A. Miles, J. Kirkham, D. Stansby, D. Papadopoulos Orfanos, J. Hamman, D. Bennett, M. Bussonnier, J. Moore, T. Augspurger, D. Cherian, M. Jones, N. Rzepka, H. Spitz, S. Verma, J. Bourbeau, A. Fulton, R. Abernathey, G. Lee, M. R. B. Kristensen and M. Durant. zarr-developers/zarr-python: v3*.1.5*. Version 3.1.5. 2025. doi: 10.5281/zenodo.17672242.

[20] Dask Development Team. Dask: Library for dynamic task scheduling. 2016.

[21] M. Rocklin. ‘Dask: Parallel Computation with Blocked algorithms and Task Scheduling’. In: Proceedings of the 14th Python in Science Conference. Ed. by K. Huff and J. Bergstra. 2015, pp. 130–136.

[22] K. J. Gorgolewski et al. ‘The brain imaging data structure, a format for organizing and describing outputs of neuroimaging experiments’. In: Scientific Data 3.1 (2016), p. 160044. doi: 10.1038/sdata.2016.44.

[23] C. R. Harris et al. ‘Array programming with NumPy’. In: Nature 585.7825 (Sept. 2020), pp. 357–362. doi: 10.1038/s41586-020-2649-2.

[24] F. Musacchio. OMIO: A policy-driven Python library for reproducible microscopy image I/O. Version v0.3.0. 2025. doi: 10.5281/zenodo.22029020.

[25] F. Musacchio. OMIO Example Datasets: Dummy Microscopy Files Plus Curated Third-Party Samples for I/O Testing. Zenodo, 2025. doi: 10.5281/zenodo.18078231.

[26] S. Crux, M. D. Roggan, S. Poll, F. C. Nebeling, J. Schiweck, M. Mittag, F. Musacchio, J. Steffen, K. M. Wolff, A. Baral, W. Witke, C. Gurniak, F. Bradke and M. Fuhrmann. ‘Deficiency of actin depolymerizing factors ADF/Cfl1 in microglia decreases motility and impairs memory’. In: bioRxiv (2024). doi: 10.1101/2024.09.27.615114. eprint: https://www.biorxiv.org/content/early/2024/09/29/2024.09.27.615114.full.pdf.

[27] F. C. Nebeling, F. Fuhrmann, M. Mittag, F. Musacchio, H. Antony, N. Gockel, L. L. Friker, S. Leonardelli, S. Filser, D. A., M. Stork, D. Bano, T. Pietsch, F. A. Giordano, Q. Zhou, S. Parrinello, M. Hölzel, U. Herrlinger, P. Salomoni and M. Fuhrmann. ‘Microglia-glioblastoma crosstalk mediates glioblastoma invasion at the far infiltration zone’. In: Immunity 59.4 (2026), 1075–1091.e4. doi: 10.1016/j.immuni.2026.03.010.

[28] F. Fuhrmann, F. C. Nebeling, F. Musacchio, M. Mittag, S. Poll, M. Müller, E. A. Giovannetti, M. Maibach, B. Schaffran, E. Burnside, I. C. W. Chan, A. S. Lagurin, N. Reichenbach, S. Kaushalya, H. Fried, S. Linden, G. C. Petzold, G. Tavosanis, F. Bradke and M. Fuhrmann. ‘Three-photon in vivo imaging of neurons and glia in the medial prefrontal cortex with sub-cellular resolution’. In: Communications Biology 8.1 (2025). Fuhrmann, Nebeling, and Musacchio contributed equally to this work., p. 795. doi: 10.1038/s42003-025-08079-8.

[29] N. Gockel, N. Nieves-Rivera, M. Druart, K. Jaako, F. Fuhrmann, F. Musacchio, H. Antony, M. Mittag, S. Crux-Daseking, S. Poll, B. Jansone, M. Fuhrmann and C. Le Magueresse. ‘Schizophrenia-associated complement C4 impairs synaptic connectivity and decreases microglia-synapse interactions through CR3 signaling’. In: Cell Reports 45.4 (2026), p. 117161. doi: 10.1016/j.celrep.2026.117161.

[30] N. Gockel, N. Nieves-Rivera, M. Druart, K. Jaako, F. Fuhrmann, R. Rožkalne, F. Musacchio, S. Poll, B. Jansone, M. Fuhrmann and C. L. Magueresse. Example Datasets for Microglial Motility Analysis Using the MotilA Pipeline. [Data set]. 2025. doi: 10.5281/zenodo.15061566.

[31] F. Musacchio, S. Crux, F. Nebeling, N. Gockel, F. Fuhrmann and M. Fuhrmann. ‘MotilA – A Python pipeline for the analysis of microglial fine process motility in 3D time-lapse multiphoton microscopy data’. In: Journal of Open Source Software 10.116 (2025), p. 9267. doi: 10.21105/joss.09267.

[32] F. Musacchio and M. Fuhrmann. ‘ZenReg: A modular Python platform for fast and memory-efficient N-dimensional microscopy image registration’. In: (2026). Preprint. doi: 10.64898/2026.08.07.743572.

[33] F. Musacchio and M. Fuhrmann. ‘Spectral Unmixing: A modular and reproducible Python package for directed and blind spectral unmixing in multidimensional microscopy stacks’. In: (2026). Preprint. doi: 10.64898/2026.07.06.736825.

[34] F. Musacchio, H. Antony, A. Baijal, D. M. Hoffmann, F. C. Nebeling, S. Crux and M. Fuhrmann. ‘CellColoc: A modular, open-source workflow for cell colocalization, segmentation, and feature extraction in microscopy images’. In: (2026). Preprint. doi: 10.64898/2026.07.26.740771.

